# Benchmarking machine learning across development reveals stage-specific neuroanatomical sex signatures

**DOI:** 10.64898/2026.11.24.768941

**Authors:** Enerson (Livi) Poon, Hamed Honari, Javad S. Ansarifar, David S. Hong

## Abstract

Machine learning (ML) offers powerful approaches for identifying multivariate patterns in neuroimaging data, yet the biological validity and interpretability of the discriminative patterns learned by such models remain incompletely understood. Here, we benchmarked multiple ML approaches for sex classification across development (8-21 years) on structural neuroimaging data and investigated the effects of age on model performance and feature stability. Linear and nonlinear models demonstrated comparable classification performance, indicating that high predictive accuracy can be achieved without relying on highly complex architectures; however, age-stratified analyses and cross-age generalization tests demonstrated that developmental stage influences classification performance, and consistency of learned neuroanatomical features. To assess biological interpretability, we applied SHAP and LIME explainability methods and developed a comprehensive framework for translating feature attributions into biologically interpretable patterns. Sexdiscriminative regions were distributed across multiple cortical and subcortical structures and exhibited marked developmental variation. Together, these findings demonstrate the utility of interpretable ML for developmental neuroimaging and highlight the importance of evaluating predictive performance while simultaneously taking biological validity, developmental specificity, and attribution stability into consideration.

## 1. Introduction

The advent of large population-level neuroimaging datasets allows implementation of machine learning (ML) approaches to decompose complex, highdimensional brain signals in ways conventional univariate statistical methods cannot. Despite significant advances, a fundamental question remains: to what extent do these data-driven models capture genuine biological signal rather than sample-specific artifacts(Haufe et al. (2014); Woo et al. (2017))?

This question is particularly pressing in developmental neuroimaging. Brain maturation is nonlinear, region-specific and modeled by complex, interactive biological influences, making ML espectially well-suited for precise characterization of age-related neuroanatomical features. As such, flexible models, such as ensemble decision trees and deep neural networks (DNN) may capture structures missed by standard linear models. Regularization and feature-selection strategies further allow these models to accommodate many correlated, high-dimensional measures typical of pediatric neuroimaging studies. However, determining biological validity of ML in these contexts requires a framework in which the expected biology is well characterized.

Age and sex effects across development offer an ideal paradigm in pediatric populations. Specifically, age directly shapes brain maturation throughout development: cortical gray volume increases 150% in the first two years of life (Knickmeyer et al. (2008)), followed by linear white matter growth, while gray matter development follows a nonlinear, inverted-U trajectory that peaks and then declines across middle childhood and adolescence. These patterns are region-specific: frontal and parietal cortices peak in volume around age 11-12 years, and temporal regions 16 years (Gogtay et al. (2004); Mills et al. (2016); Bethlehem et al. (2022)). Sex as a biological variable is similarly well-studied across physiological and cognitive domains (Talonen et al. (2017); Trainor and Falkner (2021); Burton et al. (2023); DeCasien et al. (2026)), but how to properly interpret results are often debated. Some view sex as a strong, robust, and highly discriminant variable, while others argue that within-sex variation overshadows between-sex variation, producing a brain ‘mosaic’ of sex-related features rather than two truly distinct profiles (Joel et al. (2015)). However, recent ML studies report high sex-classification accuracy from brain imaging across modalities and methods (87–96%) (Adeli et al. (2020); Sepehrband et al. (2018); Kim et al. (2022); Ritchie et al. (2018a); Matte Bon et al. (2025a). This apparent contradiction is not surprising, as many individually weak univariate signals can combine to yield strong multivariate classification accuracy without any single strong dimorphic feature. Indeed, multivariate profiles of brain activation, brain volume, and behavior can each classify sex accurately, yet an individual’s sex typicality in one domain is largely unrelated to that in another **?**. Moreover, much of this accuracy depends on global brain size, with classification from regional volume decreasing from 86% to 65% once total volume is controlled. These observations underscore the value of multivariate ML approaches for sex-specific neuroimaging analyses and the need to evaluate what such models actually learn, as investigated here.

It is important to investigate sex and age jointly, as sex steroids link the two across time-locked developmental periods, initially through early ‘organizational’ effects that remain relatively stable through pre-adolescence (Sisk and Foster (2004); Sisk and Zehr (2005a); Koolschijn et al. (2014); Konrad et al. (2013)), followed by highly dynamic ‘activational’ changes emerging at the onset of puberty (Bramen et al. (2012); Kaczkurkin et al. (2018); Herting et al. (2015)). Sex differences in clinical phenotypes emerge in the adolescent period alongside corresponding neural endophenotypes (Kaczkurkin et al. (2018); Gobinath et al. (2015); Egloff et al. (2018); Brie et al. (2019)). For example, males are more likely to be diagnosed with schizophrenia (Womer et al. (2016)) and autism (Pierce et al. (2019)), while females are twice as likely to be diagnosed with depression starting in adolescence (Cyranowski et al. (2000); Bangasser and Wicks (2017); Breslau et al. (2017)). This suggests deep learning may better capture developmental windows of structural brain change compared to linear models, while rigorous comparison of ML models conversely allows validation of these approaches against biological frameworks comprehensively delineated in prior literature.

Existing studies of age-related cortical change present four related limitations. First, most focus on narrow, fixed-age cohorts, treating adolescence alone (Schlaepfer et al. (1995); Adeli et al. (2020), or the entire <18 population, as a single, undifferentiated cohort, obscuring substantial nonlinear change occurring across these epochs. Without distinguishing middle childhood, adolescence, and young adulthood, ML models trained broadly without considering age (Ho et al. (2020)) risk omitted variable bias, treating neurodevelopmental trajectories as a latent confound systematically correlated with both predictor and outcome, yet excluded from the feature space, potentially masking how age-related maturation modulates the trajectory of sexual dimorphism in neuroanatomy. Second, prior work applying single ML architectures to sex classification in developmental neuroimaging samples in isolation (Sepehrband et al. (2018); Adeli et al. (2020); Matte Bon et al. (2025b)), and explored explainability separately (Adeli et al. (2020); Niklason et al. (2022)). Third, ML approaches often draw conclusions solely from patterns present in the training data, weakening our confidence that a model’s predictions reflect genuine biological mechanisms (Selby et al. (2025); Novakovsky et al. (2023); Ching et al. (2018)). Finally, no study has directly compared classification consistency and feature attribution across multiple ML architectures within stratified age windows using a cross-age generalization framework.

Here, we directly address these prior gaps by identifying sex-specific regions (SSRs) across developmental stages, providing one of the first direct comparisons of standard ML approaches: Linear Support Vector Machine (LSVM), which performs maximal-margin separation in high-dimensional space; Random Forest (RF), an ensemble tree-based model with built-in feature selection; Extreme Gradient Boosing (XGBoost), a state-of-the-art gradient-boosting implementation; and a deep neural network (DNN), which learns hierarchical, nonlinear feature representations through successive layers of transformation. We benchmark all against logistic regression (LOG) as an interpretable linear baseline, guarding against algorithm-specific biases and enabling cross-validation of feature-importance patterns. We further stratified our training data into biologically distinct age bins overlapping with Pre-Adolescence (6–11 years), Adolescence (12–16 years), and PostAdolescence(17–21 years) thus implementing a cross-age generalization approach. By analyzing cross-age generalization performance and feature importance across age strata, we can distinguish sex differences that are stable and pre-existing from those that emerge or reorganize during pubertal maturation.

Finally, because ML approaches are often impeded by a “black-box” tradeoff between predictive accuracy and biological interpretability (Novakovsky et al. (2023); Selby et al. (2025); Ching et al. (2018)), we paired our five-model framework with explainable AI techniques SHapley Additive exPlanations (SHAP) and aggregated Local Interpretable Model-agnostic Explanations (LIME) to assess the robustness of the developmental SSR patterns across learning architectures.

## 2. Results

### 2.1. Accuracy of ML Approaches in Classifying Sex: Comparable Linear and Deep Learning Performance

We were motivated to rigorously examine the utility of ML models in developmental populations and examined the sensitivity of these approaches in classifying individual sex by comparing linear and ML model performance. Using processed neuroimaging data from the Philadelphia Neurodevelopmental Cohort dataset (PNC; N=1,381, ages 8-21), we benchmarked five models’ discriminative capacity via repeated *k* fold cross-validation across all ages. We found that linear (LSVM 82.3%, LOG 81.5%) and most ML classifiers performed within a few points of one another; only RF (75.6%) trailed appreciably. The DNN achieved the highest discrimination with AUC (0.896). Linear (LSVM, LOG) and deep learning (DNN) models were statistically comparable given overlapping fold-wise standard deviations.

Because accuracy is not bottlenecked by model capacity, this convergence matters more for interpretation than for benchmarking across otherwise dissimilar ML algorithms. Features fed to a well-regularized network are likely to represent true properties of the underlying neuroanatomical signal rather than idiosyncrasies of any particular estimator. Sex-specific signal for features that are uncorrected for global volume are strong and largely low-order: global brain size separates the sexes with an effect size around Cohen’s *d* 1.2-1.5. Here we used volume-corrected metrics which still performed well across models, indicating a relationship that a regularized linear decision boundary captures almost in full. When the Bayes’ most optimal boundary is close to linear, flexible learners can at best match a linear function, and their extra capacity is spent modeling noise rather than signal. This is consistent with tabular-learning literature, where well-tuned linear models and gradient-boosted trees remain highly competitive with deep networks on standardized, moderately-sized tabular data. The feature-to-sample ratio (225 features; *N* = 1,381 pooled, and as few as 327 per stratum) further favors strongly regularized models. Rather than causing the deep and boosted models to fail, this regime narrows the gap with linear solutions: with per-cohort architectures, dropout (0.3-0.5) and weight decay **(Table of hyperparameters, Methods)**, the DNN was regularized enough to recover residual nonlinear structure, which is reflected in its leading AUC.

### 2.2. Theory-Driven Age Stratification Impact on Model Performance for Sex Classification

To more closely evaluate developmental age effects, we compared model performance *within* three age strata approximating biological transitions across adolescence: Pre-Adolescents (8–11 years), Adolescents (12–16 years), and Post-Adolescents (17–21 years) (Table 1b). Within-cohort accuracy increased for every model with each developmental stage, rising from 67.972.5% in Pre-Adolescents to 77.0-84.1% in Post-Adolescents, with the DNN performing best in the youngest (72.5%) and oldest (84.1%) cohorts, indicating that the same anatomical pattern becomes progressively more separable across puberty, rather than being replaced.

We also observed that the pooled benchmark exceeded within-cohort accuracy for Pre-Adolescent and Adolescent groups. The Post-Adolescent within-cohort accuracy surpassed the pooled benchmark, both for LOG (82.8%) and DNN (84.1%), consistent with existing biological frameworks in which sex-specific differences emerge during the transitional adolescent period, resulting in more distinct profiles in adulthood. The pooled score benchmark thus putatively averages a strong Post-Adolescent signal with a weaker PreAdolescent one, yielding intermediate prediction performance (Section 1b).

### 2.3. Examination of Age using a Cross-age Generalization Approach and Consistency of Model Performance

Next, to assess how classic ML models manage non-age-labeled data, we conducted a cross-age generalization analysis, where models trained exclusively on one developmental stratum were tested on the two held-out age strata. We designed this classification paradigm to examine if age must be considered to produce accurate sex classification, as well as the impact of age on various ML classification accuracies on sex.

All cross age generalization directions remained well above chance; logistic regression generalized best in three of six directions and DNN in two, including the strongest single generalization (Adolescent Post-Adolescent, 80.4%, AUC 0.890). Generalization onto the Pre-Adolescent cohort was consistently the weakest, mirroring its lower within-cohort separability.

### 2.4. Biological Interpretability and ML Explainers

Because the deep and linear models achieved comparable accuracy and AUC, model choice here is better guided by the intended use of the model than by predictive performance. For binary sex classification from structural ROI features at this sample size, the relevant trade-off is in the type of interpretability each approach affords. Deep models can recover nonlinear feature interactions, which post-hoc attribution methods (such as SHAP & LIME) can then summarize; linear models instead offer direct, coefficient-level transparency, which is often preferred in settings such as epidemiology where effect estimates must be reported and scrutinized.

While DNNs enable accurate classification of complex brain imaging phenotypes, such as biological sex, their inherent opacity limits scientific explainability and clinical translation. Explainable artificial intelligence (XAI) methods are essential to align ML outputs with established neurobiological knowledge and thereby build trust between computational and clinical audiences (Holzinger et al. (2019, 2022)). As we showed that neither DNN nor linear models are inferior in accuracy, the choice of nonlinear structure or coefficient interpretability is undetermined. Thus, we evaluated this premise against our deep learning models’ explainability advantage over linear approaches.

Here, we applied two complementary methods: SHAP (Lundberg and Lee (2017)), a game-theoretic attribution allocating a prediction among features according to their marginal contributions, and cohort-aggregated LIME (Ribeiro et al. (2016)). For each cohort, we summarized each region’s mean absolute contribution and assessed reproducibility across cross-validation folds and concordance between methods.

Because single-model feature rankings are unstable at this feature-to-sample ratio, we required reported regions to be reproducible across cross-validation folds, rather than interpreting any single run. Three candidate feature parameterizations were considered *a priori* : FreeSurfer-derived volumes, cortical thickness and surface area together, and all three features jointly. The primary analysis used subcortical volumes with cortical thickness and area, excluding regional cortical volume, which is closely approximated by the thickness-area product (Winkler et al. (2018)) and hence including it alongside adds no independent information but rather introduces collinearity. SHAP summary beeswarm plots for thickness, volume, and area feature sets respectively (Supplemental figures 18, 14) and all features (Fig. 4) show the top 10 important features per group (top 20 important features are reported as exploratory data analyses). We further examined reduced feature sets to analyze univariate patterns and feature interactions using the CLE-SH package (Lee et al. (2026b)), 18 to 4 **??**. In the SHAP beeswarm plots, each point represents one participant-feature pair. Its horizontal position provides the SHAP value, the change in the model’s predicted probability of the female class attributable to that feature for that participant. Positive values push the prediction toward female while negative toward male.

To independently interpret our findings, we implemented a parallel modelagnostic gnidivor approach, cohort-aggregated LIME(Ribeiro et al. (2016)), which leverages interpretable models to provide localized explanations based on linear approximations arpound each prediction. Aggregated LIME, by contrast, reduces participant-level importance to a single value per region: the mean signed weight of the local linear surrogates, averaged over all explained participants. Positive weights (blue) indicate that higher regional values push the prediction toward female, and negative weights (red) toward male. In aggregated LIME, regions are ranked by mean absolute attribution, and a highly ranked region with a short signed bar indicates strong but direction-heterogeneous contributions.

**Table 1:** Model Performance; panels (a–c) show the model performance comparison for different scopes of the analysis. Panel (a): Pooled sample benchmark: sex-classification performance on the full cohort (*N* = 1,381, ages 8-21) using 10-fold stratified cross-validation (mean s.d. across folds). 201 subjects were fit within each training fold only. F1-Score is the harmonic mean of precision and recall; AUC (Area Under the ROC Curve) quantifies class separability, with higher values indicating better discrimination. Bold denotes the best value per column. Panel (b): Within-age cross-sectional performance (5-fold stratified cross-validation, mean across folds) for each model trained and tested within the same developmental stratum. Bold values indicate the best performance in each column. Classification accuracy increased monotonically with developmental stage for all models (pre-adolescent *<* adolescent *<* post-adolescent), a pattern indicative of progressively intensifying neuroanatomical sex differences across the pubertal transition. Pre-adolescent (*n* = 327), Adolescent (*n* = 589), Post-adolescent (*n* = 465). Panel (c): Cross-age generalization performance: each model was trained on one developmental stratum and tested on another (train test; Pre = Pre-Adolescent, Adol = Adolescent, Post = Post-Adolescent). Cells report test accuracy (%) with AUC in parentheses; bold denotes the best model per generalization direction. All cross age generalizations remain well above chance (50%), indicating a shared, developmentally stable sex-discriminative signal; cross age generalization is asymmetric, with generalization onto the pre-adolescent cohort consistently the weakest.

| **Model** | **Accuracy** | **F1 Score** | **AUC** |  |  |  |
|---|---|---|---|---|---|---|
| LSVM | **82.3%** ± 2.6 | **0.823** ± 0.026 | 0.887 ± 0.021 |  |  |  |
| DNN | 81.5% ± 1.5 | 0.815 ± 0.015 | **0.896** ± 0.019 |  |  |  |
| LOG | 81.5% ± 1.9 | 0.815 ± 0.019 | 0.895 ± 0.021 |  |  |  |
| XGB | 78.6% ± 3.3 | 0.785 ± 0.033 | 0.869 ± 0.032 |  |  |  |
| RF | 75.6% ± 4.4 | 0.756 ± 0.044 | 0.836 ± 0.042 |  |  |  |
|  | **Pre-Adolescent** |  | **Adolescent** |  | **Post-Adolescent** |  |
| **Model** | Accuracy | AUC | Accuracy | AUC | Accuracy | AUC |
| LOG | 71.0% | 0.776 | **80.0%** | **0.883** | 82.8% | **0.914** |
| LSVM | 69.1% | 0.748 | 77.3% | 0.864 | 81.3% | 0.897 |
| RF | 68.5% | 0.757 | 76.6% | 0.842 | 77.0% | 0.852 |
| XGB | 67.9% | 0.756 | 77.8% | 0.863 | 79.6% | 0.868 |
| DNN | **72.5%** | **0.788** | 77.4% | 0.878 | **84.1%** | 0.901 |
| **Model** | Pre→Adol | Pre→Post | Adol→Pre | Adol→Post | Post→Pre | Post→Adol |
| LOG | 75.4 (.812) | 73.1 (.807) | **76.2 (.814)** | 79.6 (.885) | **71.9 (.775)** | **77.1 (.859)** |
| LSVM | 72.0 (.778) | 71.4 (.788) | 74.9 (.811) | 78.7 (.882) | 68.8 (.755) | 74.9 (.836) |
| RF | 76.2 (.843) | 77.4 (.839) | 68.8 (.766) | 76.3 (.852) | 70.0 (.754) | 75.6 (.840) |
| XGB | 76.4 (.827) | **77.9 (.844)** | 70.6 (.774) | 79.8 (.880) | 70.0 (.742) | 76.1 (.856) |
| DNN | **78.4 (.855)** | 76.3 (.854) | 74.6 (.815) | **80.4 (.890)** | 71.6 (.785) | 76.2 (.861) |

**Figure 1:**
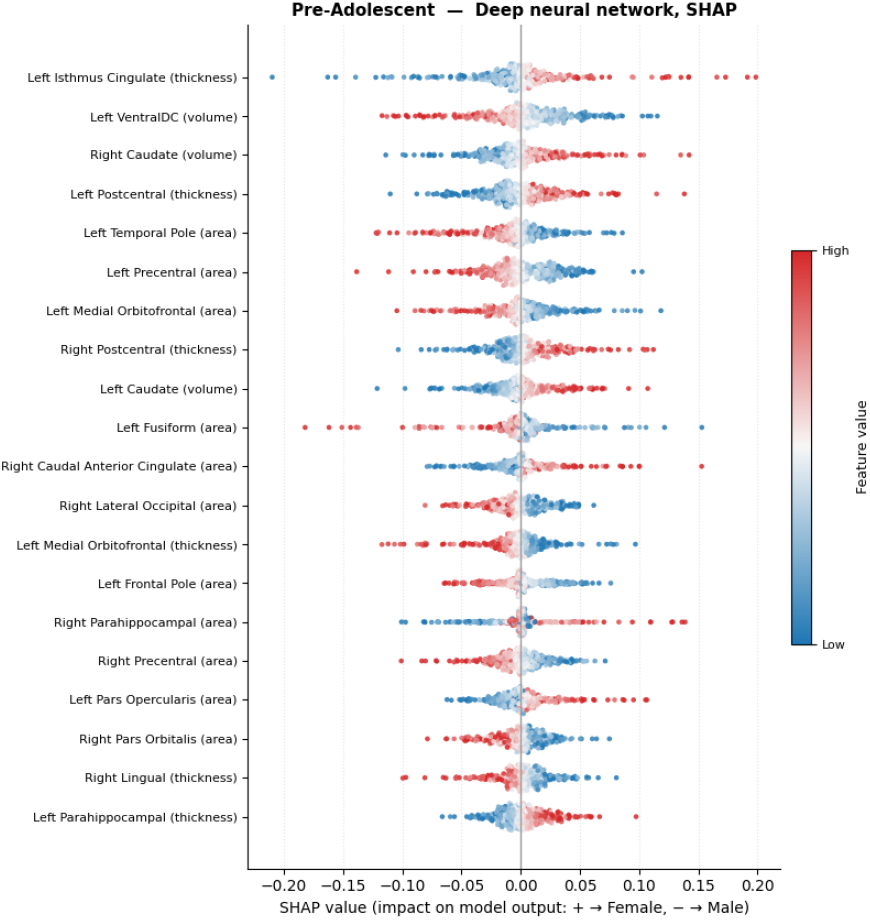
Panel (a). Group 1: Pre-Adolescent

**Figure 2:**
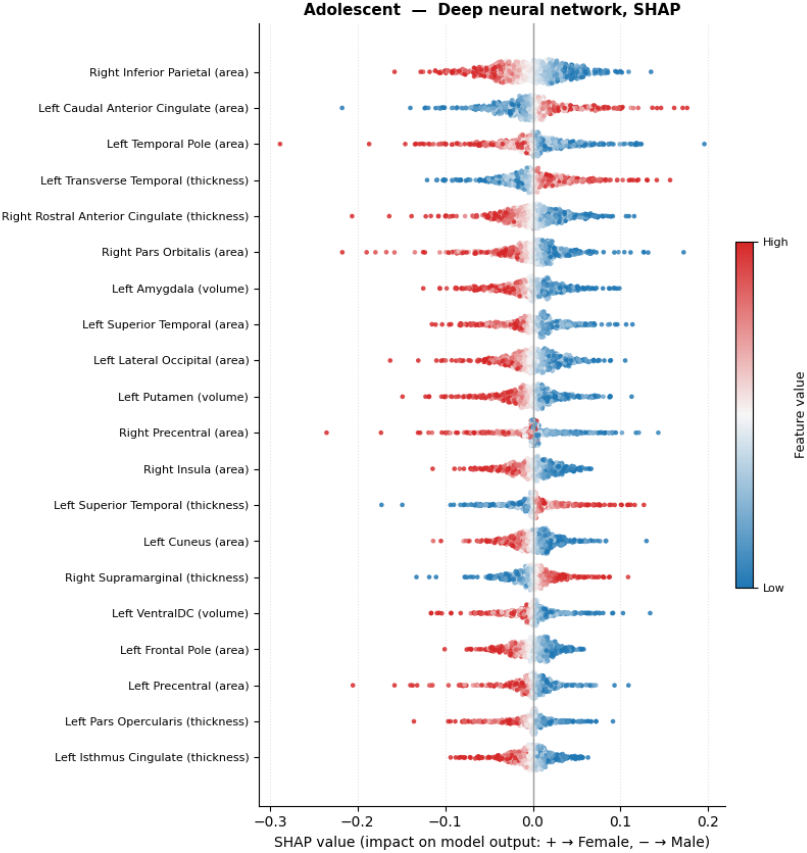
Panel (b) Group 2: Adolescent

**Figure 3:**
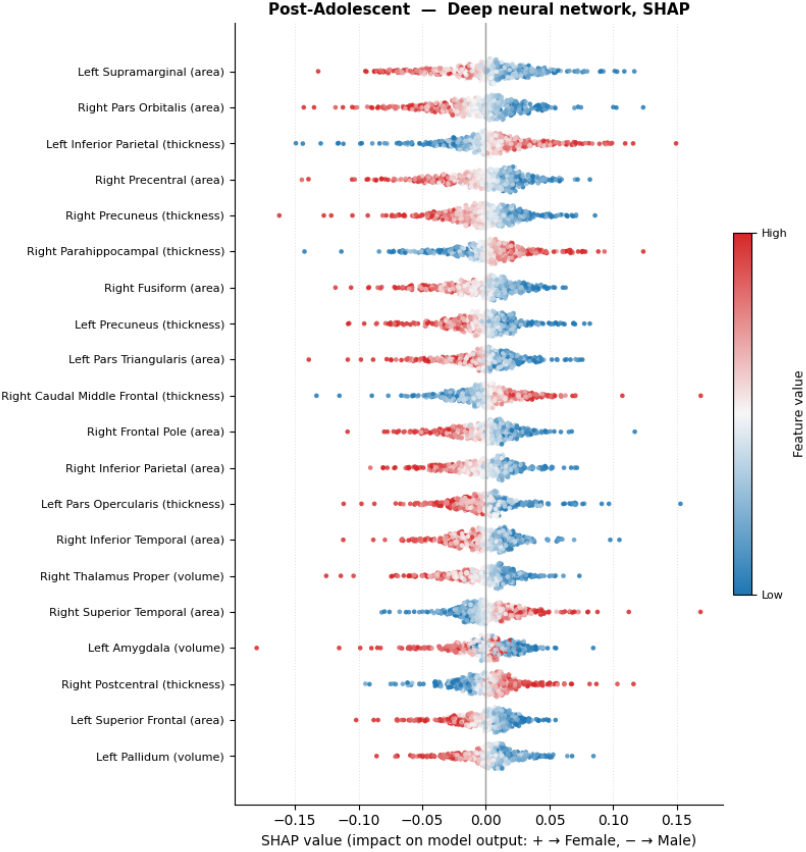
Panel (c) Group 3: Post-adolescent

**Figure 4:** Cross validated SHAP explanations of the deep neural network sex classifier within each developmental cohort. Beeswarm plots of SHAP values for (a) the Pre-adolescent, (b) Adolescent, and (c) Post-adolescent within-cohort analyses. Within each analysis, SHAP values were pooled across the five outer cross-validation folds: each dot represents one participant feature pair. Panels show the top 20 features with the largest pooled mean absolute SHAP value in descending order. Positive values push the prediction towards Female, negative values toward Male, and each dot encodes the subject’s feature value.

**Figure 5:**
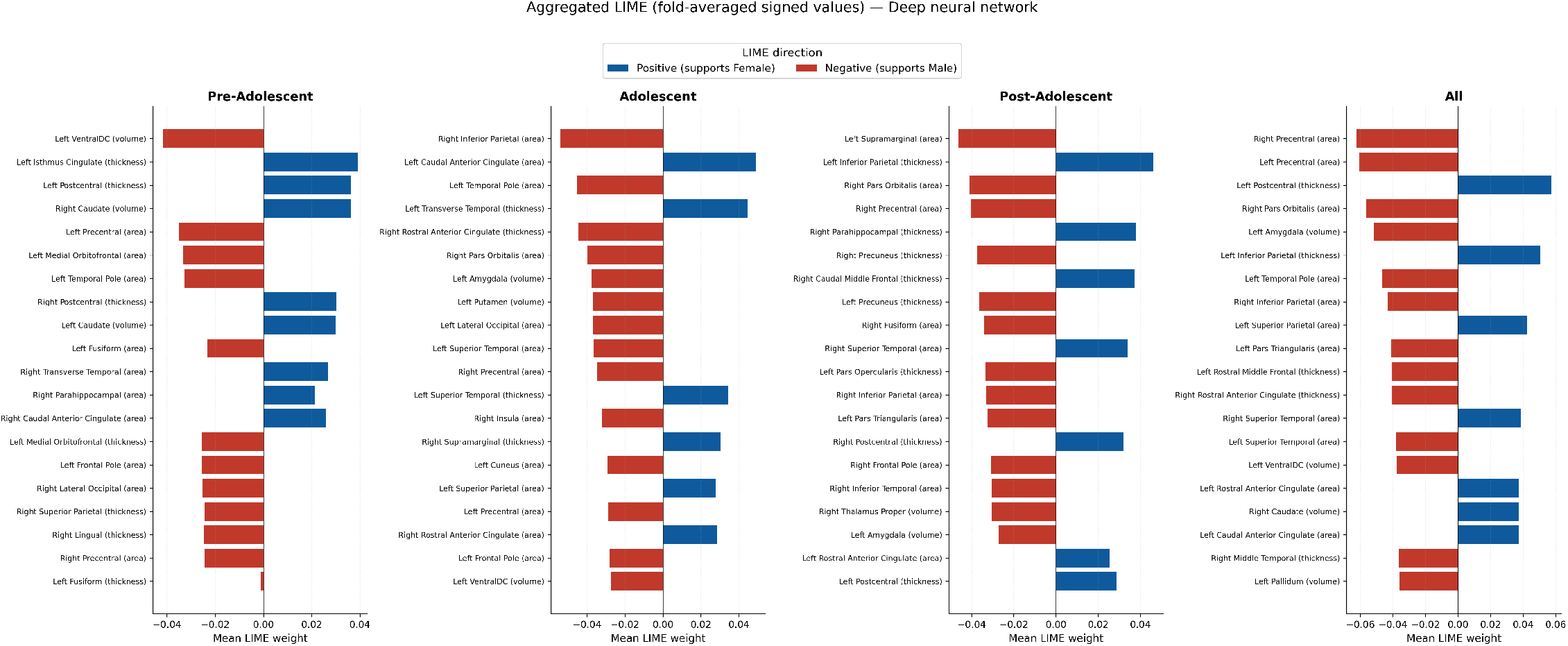
Aggregated LIME feature attributions for the three age cohorts, and pooled cohort (all) computed from the corresponding per-cohort deep neural network classifiers. Each bar is the mean signed LIME weight of a brain region across all subjects in the cohort; positive values indicate features that pushed the classifier toward a female prediction and negative values toward a male prediction. Aggregated LIME closely matched the SHAP attributions in every cohort (Spearman *ρ* 0.99), providing cross-method confirmation of the sex-discriminative regions identified by SHAP.

### 2.5. Sex-Discriminative Regions and Their Developmental Trajectory

SHAP and cohort-aggregated LIME are methodologically distinct, the former deriving from a game-theoretic allocation of a prediction among features and the latter from an average of local linear surrogates. We found the two methods’ importance profiles corresponded closely within each age cohort (Spearman *ρ* = 0.987, 0.993 and 0.992 for the Pre-Adolescent, Adolescent and Post-Adolescent cohorts (top-20 overlap 17/20, 18/20 and 18/20), approximately twelve standard deviations above the permutation null for 157 features (null mean *ρ* 0, SD = 0.080). Kendall’s *τ* behaved similarly (0.907, 0.932, 0.926). We also found cross-method agreement substantially exceeded either method’s own fold-to-fold reproducibility. The correlation between a single fold’s ranking and the fold-averaged ranking of the same method was only *ρ* = 0.44 0.75 (median 0.66) for SHAP and 0.44-0.75 (median 0.67) for aggregated LIME, far below the *ρ* = 0.98 observed between the two different methods.

Aggregated LIME tracked mean absolute SHAP almost proportionally (coefficient of variation of their ratio 0.07–0.08; *r* = 0.989 on raw and 0.988 log scales), consistent with the fact that Kernel SHAP and a continuous locallinear surrogate are both local additive approximations of the same trained network, fitted around the same instances, rather than independent probes. We treat their agreement as evidence of internal consistency and correct implementation, not as independent methodological validation, which requires a method that does not rely on local linearity (permutation or ablation importance). Confidence in the reported regions therefore rests primarily on their stability across folds (Fig. 6).

Chance-corrected indices were modest, yet substantial for each cohort: Cohen’s *κ* for top-20 membership was 0.83 (Pre-Adolescent), 0.89 (Adolescent) and 0.89 (Post-Adolescent), the quadratically weighted *κ* on importance deciles was 0.98 in all three cohorts, and rank-biased overlap was 0.81, 0.97 and 0.89. Bootstrap confidence intervals overlapped substantially across cohorts, which is expected as only 20 features enter the comparison. Given that single-fold rankings were unstable at this feature-to-sample ratio (157 features; *n* = 327-589 per cohort), we treat cross-fold stability as the primary basis for confidence: a region was only retained if it ranked in the top 20 in at least 4 of 5 folds independently for both SHAP and LIME.

Our importance profiles were largely specific to developmental stage compared to the strong cross-method agreement observed within each cohort. Rank correlations between cohort profiles, computed across all 157 features, were low (SHAP *ρ* = 0.21, 0.01 and 0.14 for Pre-Adolescent-Adolescent, Pre-Adolescent-Post-Adolescent and Adolescent-Post-Adolescent; aggregated LIME 0.21, 0.01 and 0.12). These values fall well below each method’s own fold-to-fold reproducibility (median *ρ* = 0.66–0.67), indicating that between-cohort differences exceed the variability attributable to attribution estimation and therefore reflect genuine differences in what each cohort’s models learned. Conservation among the leading regions declined monotonically with developmental distance: the Pre-Adolescent and Adolescent top-20 sets shared 7 of 20 features (chance expectation 2.5; hypergeometric *p* = 0.005), the Adolescent and Post-Adolescent sets 5 of 20 (*p* = 0.09), and the two most distant cohorts only 3 of 20, no more than expected by chance (*p* = 0.49). Only two features, right precentral and right pars orbitalis surface area, remained in the top 20 at every stage (Table 3). Moreover, the features shared between adjacent cohorts differed in composition: the PreAdolescent–Adolescent overlap comprised the early subcortical and polar signature (ventral diencephalon, temporal pole), whereas the Adolescent-PostAdolescent overlap comprised the emerging limbic-parietal signature (amygdala, inferior parietal cortex), consistent with progressive shifting of the discriminative set rather than fluctuation around a fixed architecture. Taken together, these analyses indicate stage-dependent successive reorganization of the sex-discriminative feature space around a conserved core, consistent with a two-stage organizational-activational model. Indeed, the stage-specific signatures followed established biological expectations in Pre-Adolescents, the most stable sex-discriminative features were the ventral diencephalon (which includes the hypothalamus) and caudate. In Adolescents, the inferior parietal cortex, rostral anterior cingulate and left temporal pole emerged alongside pars orbitalis, amygdala and superior temporal regions. In Post-Adolescents, the stable set emerged as predominantly cortical, including inferior parietal cortices, precentral cortex, and orbitofrontal (pars orbitalis), and the parahippocampus emerging.

Our findings demonstrate progression from an early hypothalamic/striatal signature in early adolescence, a transitional adolescent limbic profile, and a later consolidation into a cortical associative/sensorimotor medial profile, recapitulating a well-established two-stage biological model in which perinatal organizational effects establish a sexually differentiated substrate that is subsequently remodeled by pubertal gonadal hormones Schulz et al. (2009). Developmental re-ordering is thus concentrated among the most discriminative regions rather than distributed across the full feature set. For all reported regions, we emphasize that signed mean attribution was small relative to the absolute attribution (median signed */* abs 0.05-0.09), the expected signature of a genuinely discriminative feature whose contribution reverses with the participant’s own value. We therefore describe these regions as sex-*discriminative* rather than uniformly larger in one sex, interpreting directionality from the SHAP dependence plots rather than from signed means.

**Table 2:** Agreement between SHAP and aggregated LIME feature-importance profiles within each developmental cohort, computed on fold-averaged mean absolute attributions (*n* = 157 features). Global indices are computed over all features; chance-corrected and top-weighted indices focus on the head of the ranking. The weighted *κ* computed on raw ranks is numerically identical to Spearman *ρ* (absolute difference *<* 10*^−^*^15^), a consequence of the uniform marginals produced by ranking 51, and is reported here only to document that redundancy; it should not be presented as independent corroboration.

| **Agreement index** | **Pre-adolescent** | **Adolescent** | **Post-adolescent** |
|---|---|---|---|
| *Global association (all 157 features)* |  |  |  |
| Spearman *ρ* | 0.987 | 0.993 | 0.992 |
| Kendall *τ* | 0.907 | 0.932 | 0.926 |
| Weighted *κ*, raw ranks | 0.987 | 0.993 | 0.992 |
| *Chance-corrected and top-weighted* |  |  |  |
| Weighted *κ*, deciles | 0.983 | 0.988 | 0.982 |
| Rank-biased overlap | 0.813 | 0.965 | 0.886 |
| Cohen *κ*, top-20 | 0.828 | 0.885 | 0.885 |
| Jaccard, top-20 | 0.739 | 0.818 | 0.818 |

## 3. DISCUSSION

In this study, we systematically evaluated ML approaches for classifying sex from structural neuroimaging data across childhood, adolescence, and young adulthood; discrete developmental periods characterized by wellestablished nonlinear patterns of brain maturation overlapping with known changes in sex steroid influence and puberty. Our analyses provide a principled assessment of whether ML methods offer a meaningful advantage over linear approaches in capturing the more complex relationships that emerge during adolescence, addressing the need for methodological rigor when applying flexible ML algorithms to data with underlying ground-truth biological frameworks.

Prediction accuracy for sex classification increased with age for both linear and deep learning models, peaking in the Post-Adolescent cohort and nominally exceeding the full, non-age-stratified pooled sample. This pattern supports a framework in which the most pronounced sex-discriminant brain features emerge after puberty. In line with existing literature, loworder sex classification proved fairly robust across both linear and nonlinear methods, although we observed incremental gains in performance for more complex, hierarchical models such as DNNs. A similar tension appears in parallel work: deep models trained directly on raw structural imaging data have been shown to yield accurate, heritable brain-age biomarkers, yet largescale comparisons across imaging phenotypes have found that simple linear models can perform on par with deep learning except at very large sample sizes (Schulz et al. (2020)).

**Table 3:**
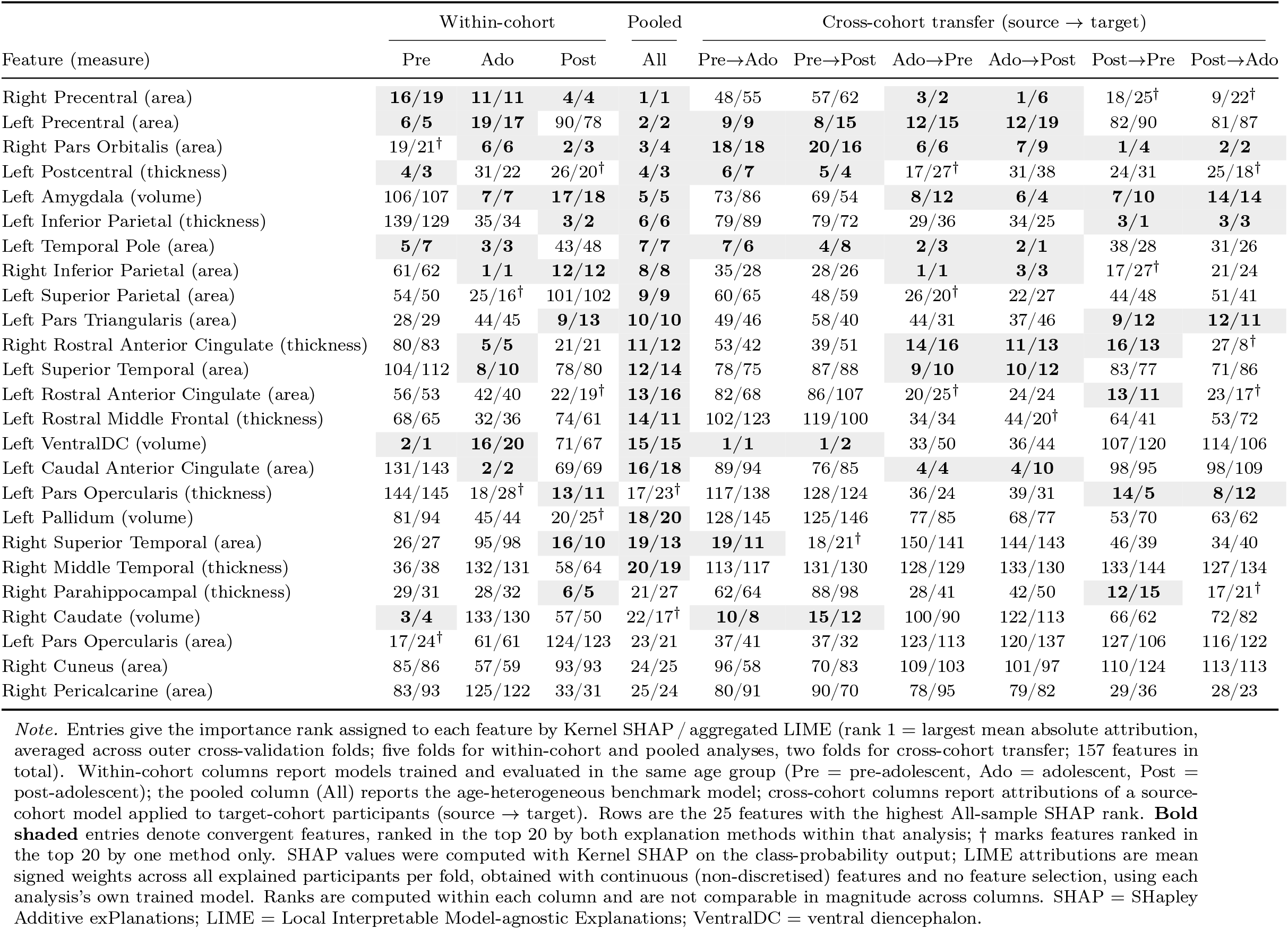
Convergence of SHAP and LIME feature-importance rankings for deep neural network sex classification within developmental cohorts, in the pooled sample, and under cross-cohort transfer.

**Figure 6:**
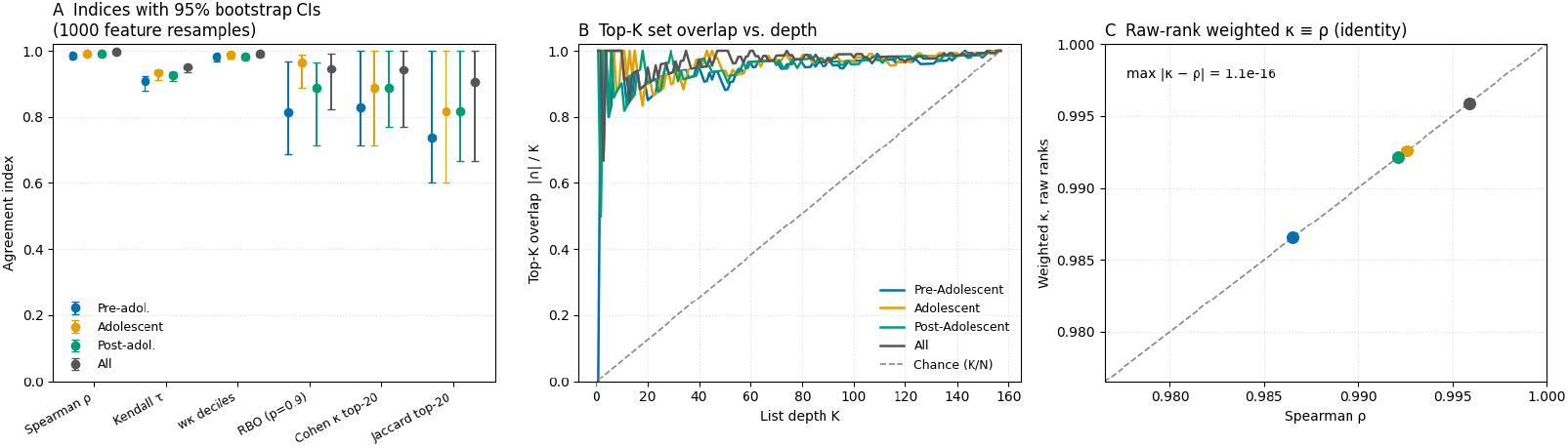
Agreement between SHAP and aggregated LIME across developmental cohorts. (A) Complementary agreement indices with 95% bootstrap confidence intervals obtained by resampling features. Global rank indices (Spearman, Kendall) are uniformly high, whereas chance-corrected and top-weighted indices are lower but still substantial, and their confidence intervals overlap across cohorts. (B) Overlap between the two methods’ top-*K* feature sets as a function of list depth *K*, with the overlap expected by chance (*K/N*) shown for reference; agreement greatly exceeds chance at every depth. (C) The quadratically weighted *κ* computed on raw ranks plotted against Spearman *ρ* for each cohort: the two coincide on the identity line to machine precision, demonstrating empirically that weighted *κ* adds no information beyond the rank correlation unless importance is first categorized.

First, our benchmarking results support the biological validity of MLbased sex classification and strikingly follow the expected biological pattern of dimorphic emergence across development. Existing frameworks hold that developmental sex differences are driven primarily by sex-steroid influence, with initial “organizational” effects imprinted during the perinatal period. These are followed by relatively minimal dimorphic separation across middle childhood (represented here in the Pre-Adolescent cohort), when circulating testosterone and estrogen profiles remain roughly equivalent between boys and girls (Blakemore (2008)). The onset of puberty, typically at age 11 years for girls and 12 years for boys (Blakemore (2008)), with its concomitant surge of testosterone in males and estrogens in females, is thought to drive divergent structural brain profiles during the “activational” period of adolescence, which then stabilizes into the patterns observed in young adulthood (here, the PostAdolescent cohort). Our within-group, age-stratified analyses supported this progression for all models: accuracy improved from 67–72% in the middlechildhood cohort to 77–84% in the oldest.

**Figure 7:**
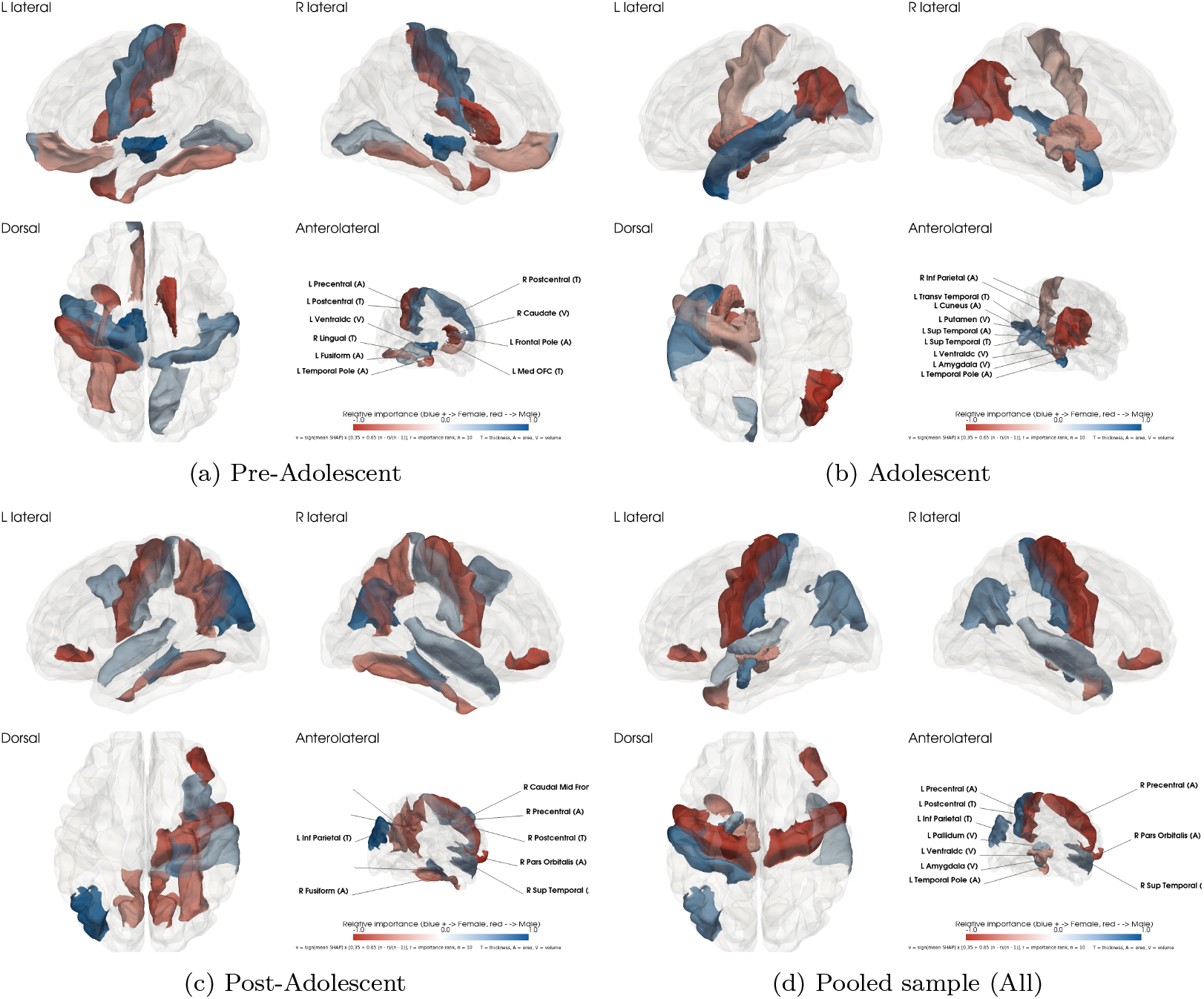
Convergent SHAP-LIME features for the deep neural network sex classifier within each developmental cohort and in the pooled sample. Each panel shows, for one analysis — (a) Pre-adolescent, (b) Adolescent, (c) Post-adolescent, (d) pooled ageheterogeneous benchmark (All). The five strongest Female-associated and five strongest Male-associated convergent features, defined as features ranked in the top 20 by both Kernel SHAP and aggregated LIME in that analysis (fold-averaged mean absolute attributions). Color hue encodes attribution direction, determined by the sign of the fold-averaged mean signed SHAP value: blue, higher feature values shift the predicted probability toward Female; red, toward Male. Color saturation encodes importance rank among the displayed features, mapped linearly from the highest rank (full saturation) to the lowest (display floor of 0.35). Table 3 provides a comprehensive list of ranking convergence for the within, benchmark, and cross analysis.

Second, our cross-age generalization paradigm shows that all models achieve prediction accuracy well above chance, but with a specific directionality. Classifiers trained on the Pre-Adolescent cohort generalized relatively well to older cohorts, indicating enduring sex-discriminative features, whereas Adolescent and Post-Adolescent classifiers generalized weakest to the Pre-Adolescent cohort reflecting its lower intrinsic separability, and suggesting a methodological pitfall when age is treated as a unitary construct. Classifiers trained on the Adolescent cohort performed nominally better than those trained on either the Preor Post-Adolescent cohort alone, possibly reflecting transitional SSRs bridging the two developmental extremes. Interestingly, RF and XGBoost showed the poorest cross-age generalization accuracy, which we cautiously attribute to their fixed, tree-based decision boundaries being less adaptable across developmental stages than the continuous decision surfaces of the DNN and LSVM. Together, these findings underscore the importance of accounting for age a priori when constructing the feature space for ML-based neuroimaging classification.

Third, our analyses extend the literature by using XAI methods to biologically validate model outputs, addressing the long-standing challenge of translating neuroimaging-derived models into interpretable, mechanistically grounded findings (Aas et al. (2021); Haufe et al. (2014); Woo et al. (2017)). As detailed above, SHAP and aggregated LIME agreed almost perfectly on global feature ranking. Gradient-based SHAP and local-linear LIME both approximate the same underlying local gradient, and their attributions here differed only by an approximately constant scaling factor. We therefore read the convergence as evidence of an internally consistent implementation rather than an independent cross-method validation a distinction that matters wherever “explainer agreement” is offered as a robustness argument. The more conservative, chance-corrected indices (0.83–0.89) provide a more realistic estimate of cross-method agreement at the head of the ranking; due to their bootstrap confidence intervals overlapping across cohorts, we do not interpret cohort differences in these indices.

SHAP and aggregated LIME further converged on a developmental result. Regional importance profiles were largely specific to developmental stage, with between-cohort similarity declining progressively with developmental distance and falling below within-cohort fold-to-fold variability, whereas only two features, precentral and pars orbitalis surface area, remained among the leading regions at every stage. Sex-discriminative neuroanatomy in this sample is therefore better described as a minimal early-established core of stable features on which a predominantly stage-specific profile is progressively remodeled across puberty, rather than as an unchanging architecture maintained throughout development. This characterization is consistent with longitudinal evidence that male and female structural trajectories are themselves dimorphic, such that regional sex differences emerge and fade at different ages in different systems Lenroot et al. (2007); Giedd et al. (2012); Kaczkurkin et al. (2019), and with recent longitudinal findings that pubertal development predicts structural maturation over and above age, in a sex-specific manner Beck et al. (2023); our stage-dependent discriminative maps are the multivariate consequence of that trajectory dimorphism. It is also a better fit to the two-stage organizational-activational framework than a static backbone would be: sex differences in brain structure are detectable before birth Paulus et al. (2026), and extensive reorganization concentrated in the pubertal window is the expected signature of activational gonadal-hormone effects acting on that organizationally shaped substrate Phoenix et al. (1959); Sisk and Zehr (2005b); Schulz et al. (2009); Drzewiecki and Juraska (2020). That the conserved core comprises surface area measures accords with the early, prenatally organized origin of cortical arealization and with evidence that sex differences load predominantly on surface area Ritchie et al. (2018b); Lee et al. (2026a).

The stage-specific components align closely with hormone-sensitive neuroanatomy. In Pre-Adolescents, the ventral diencephalon (including the hypothalamus) and the caudate are most prominent, consistent with sexual differentiation established by organizational steroid effects before puberty: adult dimorphism is greatest in regions with high developmental sex-steroid receptor density, an effect established on brain-size-adjusted volumes Goldstein et al. (2001). In Adolescents, the emergence of the amygdala and temporo-limbic cortex matches longitudinal evidence that medial temporal development is driven by pubertal stage rather than age alone, in a sex-specific manner Goddings et al. (2014); Neufang et al. (2009); Bramen et al. (2011). The adjacent-cohort overlaps are transitional in composition, since the features shared with Pre-Adolescents belong to the receding early signature while those shared with Post-Adolescents belong to the emerging limbicparietal one. This split marks adolescence as the transition between the two regimes. By Post-Adolescence, the discriminative set had consolidated onto association and sensorimotor cortex, consistent with the protracted maturation of these regions and with the broad cortical sex differences reported in adults, which persist, attenuated, after adjustment for total brain volume Gogtay et al. (2004); Ritchie et al. (2018b). Notably, regions where adult sex differences align with testicular hormone effects (dorsal parietal cortex, angular gyrus), overlap the inferior parietal features that emerged in our Adolescent and Post-Adolescent cohorts. We emphasize that stage-specificity of multivariate importance rankings does not contradict the replicability of adult univariate sex differences, nor does it adjudicate the debate over their magnitude once global size is controlled Ritchie et al. (2018b); Eliot et al. (2021); DeCasien et al. (2022); Sanchis-Segura et al. (2022); BrzezinskiRittner et al. (2025). Classifiers apportion credit among collinear features and can exploit distributional differences such as the greater male variance observed across the lifespan Wierenga et al. (2022); hence, the regions a model relies upon can reorder substantially across development even where many underlying univariate effects persist. Furthermore, this anatomical convergence is further supported at the molecular level, where single-cell transcriptomic profiling of adult cortex, targeted to regions with reproducible sex differences in gray matter volume, found sex-biased expression concentrated in fusiform cortex and present in intraparietal and insular regions, organized along the sensory-to-association axis and regulated by androgen, estrogen, and progesterone receptors and X-linked transcription factors. Several of these regions correspond to features our models relied on, including fusiform surface area and superior and inferior parietal cortex, which border the intraparietal sulcus. As such, our findings that regions identified by data-driven attribution in youth coincide with sites of hormoneand sex-chromosome-regulated gene expression in adults, further support the biological validity of the XAI-derived patterns. It also suggests our developmental maps can provide complementary insights on the timing of these molecular sex differences.

Methodologically, models trained on samples pooled across adolescence will recover an age-averaged composite profile corresponding to no single developmental stage; together with the growing methodological scrutiny of sex-classification studies Adeli et al. (2020); Sapuan et al. (2026), and evidence that volumetric sex classification falls from 86

Our cross-sectional, chronologically binned approach has several limitations. Age is an imperfect proxy for pubertal stage and the sex differences classifer in this work also likely reflects individual variation in pubertal tempo, as well as between-group effects between girls and boys who differ in timing of pubertal onset. While pubertal status was not available for further discrimination in this dataset, future machine learning studies should examine chronological age as well as pubertal status as a potentially more sensitive measure, ideally in longitudinal study designs. The stable regions we report reflect fold-robust sets rather than precise ranks, since individual feature ranks remained variable across folds. Because SHAP and aggregated LIME are both local gradient-based approximations, their agreement is simply a consistency check rather than independent validation. Nonetheless, by explicitly grounding interpretation in attribution stability and cross-method agreement, this work moves the research of sex-specific regions from single-model feature lists toward reproducible, developmentally resolved neuroanatomical findings, and offers a template for using XAI to track how sexually dimorphic brain organization is established and refined across adolescence – a period in which sex-biased neuropsychiatric conditions characteristically emerge (Kaczkurkin et al. (2018)).

We suggest that model choice for pediatric neuroimaging data should be centered around a particular scientific question, rather than treated as a question of a single ‘best’ architecture. If the goal is maximizing raw classification accuracy alone in feature spaces that are lower-dimensional, well-characterized, or sample-limited, a linear model remains defensible and often preferable for its lower variance, reduced overfitting when the sample-to-feature ratio is constrained, and minimal hyperparameter tuning needed to reach stable performance, consistent with observed convergence between linear and deep models at moderate sample sizes Schulz et al. (2009). Linear approaches are also directly interpretable in comparison to most ML models output from logistic regression can be derived analytically from its inputs, making it a genuine white-box method rather than one requiring post-hoc approximation.

Our results show that deep learning models may offer a distinct advantage as feature-space complexity grows and may reduce blind spots imposed by the fixed functional form of linear or margin-based models. This flexibility is particularly relevant given that few brain ROIs are truly independent of one another penalized linear models can arbitrarily downweight or discard biologically meaningful but collinear features during regularization, whereas a DNN’s distributed representation absorbs this redundancy natively. This advantage comes with a corresponding cost in interpretability, and the two model classes are not interchangeable on this dimension, a central tension in the broader field of translational neuroimaging research, where models must be trusted in proportion to both empirical performance and biological plausibility (Woo et al. (2017)). Logistic regression’s feature importance is defined in terms of deletion, e.g., how much accuracy degrades when a feature is removed, while SHAP-based importance for a DNN is defined locally, e.g., how much a feature shifts an individual prediction. These are conceptually distinct frameworks, not two measurements of the same underlying quantity. Our cross-method agreement analysis illustrates that SHAP and LIME can be meaningfully validated against one another within a single model, because both approximate the same local decision surface, but this does not extend across model classes, since a white-box linear explainer and a black-box deep explainer are not estimating a common target. Therefore, developing importance metrics that are genuinely comparable across model classes, rather than assuming explainability is a single, transferable standard, remains an open problem for future methodological work in this space.

## 4. Methods

### 4.1. Reference Dataset

We obtained data from the Philadelphia Neurodevelopmental Cohort (PNC) (Satterthwaite et al. (2016)), a community-based study of brain development comprising subjects aged 8-21 years and including questionnaires, a computerized neurocognitive battery, structural and functional MRI, and diffusion tensor imaging. (see Table 4 for full demographic characteristics). Of this total, 1, 601 participants (16.8%) had structural MRI scans, representing those who consented to imaging, had no contraindications, and completed the scanning protocol. We included only participants with complete and successful FreeSurfer reconstructions across all the feature categories (*n* = 1, 452 or 90.7%). This excluded scans with processing failures due to poor quality, contrast, and/or reconstruction errors. To ensure phenotype completeness, we retained participants with complete demographic data (sex, age at scan) and restricted to the first assessment to avoid repeated measures, yielding our final analytical sample of *n* = 1, 381 participants. The following table shows the demographic information (see Table **??** for full demographic characteristics) of all PNC participants.

### 4.2. Participants

The final analytic sample comprised 1,381 youth (741 females, 53.7 %; age range = 8-21 years, *M SD* = 14.42 3.45 years). To evaluate analytical approaches incorporating age and development, two approaches were used: (1) modeling the entire PNC cohort, and (2) sub-grouping participants into three cohorts to evaluate the accuracy of ML models in predicting sexspecific brain regions. As pubertal status was not available (ie: Tanner staging or testosterone/estradiol labs), groups were separated by chronological age as a proxy for pubertal development based on average onset of pubertal development (Brix et al. (2019)). This stratification resulted in three developmentally relevant age bands: *i.* Pre-Adolescents aged 8-11 years (*n* = 327; 169 females [51.7 %]; age = 9.64 1.08 years), *ii.* Adolescents aged 12-16 years (*n* = 589; 301 females [51.1 %]; age = 14.07 1.41 years), and *iii.* Post-Adolescents aged 17-21 years (*n* = 465; 271 females [58.3 %]; age = 18.22 ± 1.12 years).

**Table 4:** Group summaries for count, age, and estimated total intracranial volume (eTIV).

| **Group** | **Count** | **Mean Age (years)** | **Age SD** | **Mean eTIV** | **eTIV SD** |
|---|---|---|---|---|---|
| Dataset | 9496 | 13.73 | 3.67 | 1,533,964.73 | 173,282.30 |
| Female | 4894 | 14.08 | 3.65 | 1,450,831.06 | 155,155.44 |
| Male | 4569 | 13.35 | 3.66 | 1,630,217.94 | 140,496.65 |
| Pre-adolescent | 3075 | 9.41 | 1.11 | 1,539,312.94 | 135,673.02 |
| Post-adolescent | 2623 | 18.33 | 1.15 | 1,521,585.46 | 199,406.24 |
| Adolescent | 3792 | 14.05 | 1.41 | 1,542,044.57 | 166,263.26 |

**Table 5:** Participants demographic characteristics by age group (*N* = 1,381).

| **Age Group** | **N** | **% Total** | **Males** | **Females** | **% Female** | **Mean ± SD** | **Range** |
|---|---|---|---|---|---|---|---|
| Pre-adolescent | 327 | 23.7% | 158 | 169 | 51.7% | 9.64 ± 1.08 | 8-11 |
| Adolescent | 589 | 42.7% | 288 | 301 | 51.1% | 14.07 ± 1.41 | 12-16 |
| Post-adolescent | 465 | 33.7% | 194 | 271 | 58.3% | 18.22 ± 1.12 | 17-21 |
| **Total** | 1,381 | 100.0% | 640 | 741 | 53.7% | 14.42 ± 3.45 | 8-21 |

### 4.3. MRI Acquisition and Data Processing

Structural MRI was acquired using a Siemens 3T TIM Trio whole-body scanner (VB17 revision, Siemens software) at the Hospital of the University of Pennsylvania (Satterthwaite et al. (2016)). T1-weighted anatomical images were obtained with a magnetization-prepared rapid gradient-echo (MPRAGE) sequence: TR = 1810 *ms*, TE = 3.51 *ms*, TI = 1100 *ms*, flip angle = 9°, FOV = 180 × 240 *mm*, matrix = 192 × 256, 1 *mm* isotropic voxels (Satterthwaite et al. (2016)). The T1-weighted images were utilized from all available participants and were processed via FreeSurfer (Fischl (2012)). To account for individual variances in global brain size, brains were normalized via estimated total intracranial volume (eTIV) (Sargolzaei et al. (2015)). Then, features for all subjects were calculated: volume (*mm*^3^), surface area (*mm*^2^), and average cortical thickness (*mm*), if the region was a subcortical feature, only the volume of the structure was computed. Features were extracted using the default FreeSurfer atlas, Desikan-Killiany (Desikan et al. (2006)).

### 4.4. Feature Selection

Cortical thickness, surface area, and gray matter volume were extracted from 34 bilateral regions-of-interest (Desikan-Killiany atlas; 68 features per metric, total 204 cortical) alongside volumes of key subcortical structures (hippocampus, amygdala, thalamus, caudate, putamen, pallidum, nucleus accumbens; bilateral), corpus callosum segments, and ventral diencephalon (20 subcortical features). We further excluded artifact-prone, non-specific or summary measures (such as ventricles, CSF, hypointensities, etc.), retaining 225 biologically relevant features which focus on individual lowest-order brain regions. Of these, the primary analysis set comprised cortical thickness and surface area together with the subcortical volumes (157 features), excluding regional cortical volume for the reasons detailed below; all interpretability results in the main text derive from this set.

**Figure 8:**
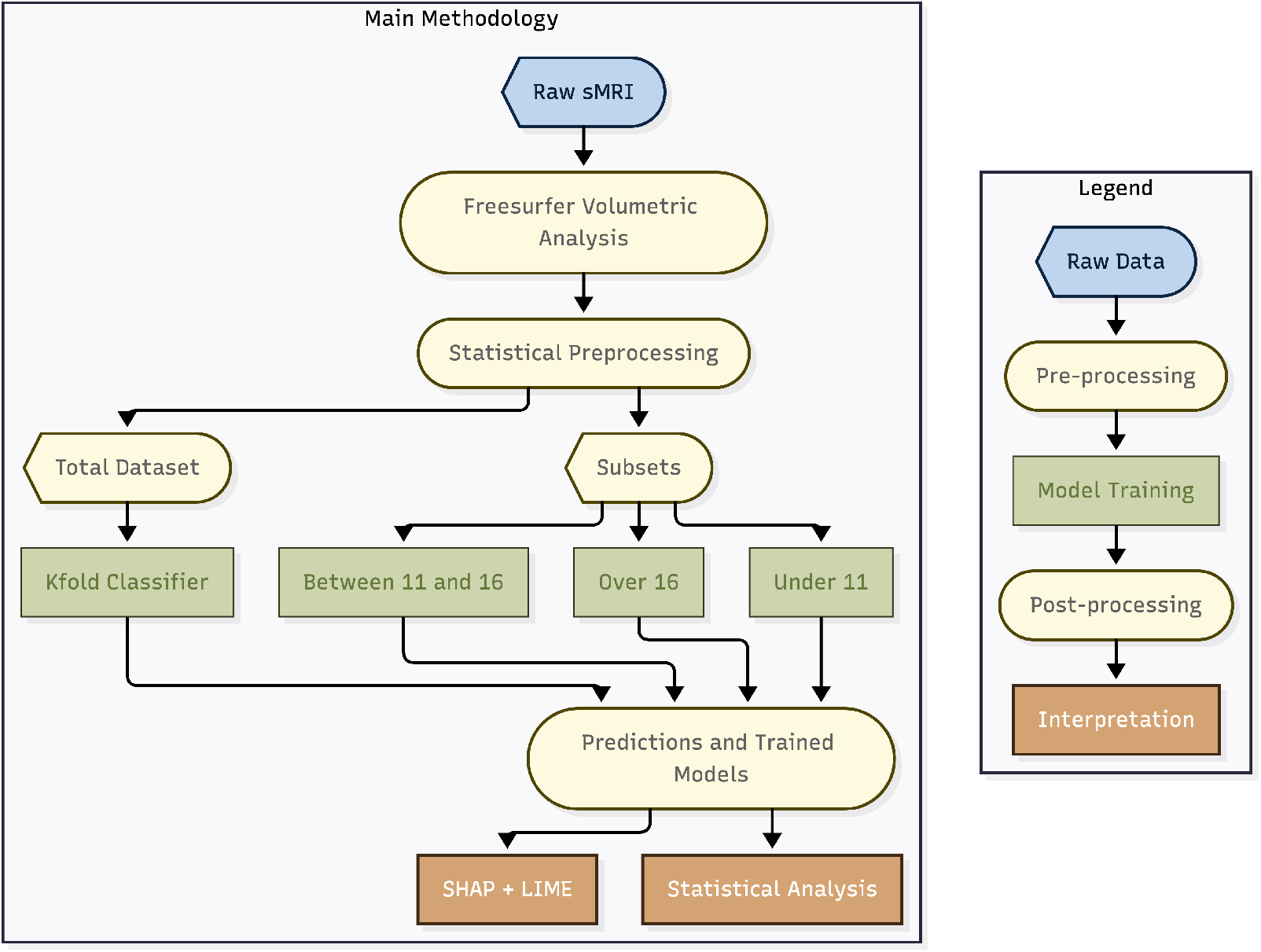
Processing Workflow: Raw MRI’s are preprocessed through FreeSurfer’s Volumetric Group Analysis algorithms to quantify the volume, thickness, and area of 35+ ROI’s. After global variables, means, and insignificant brain regions were removed, each subject’s values were normalized over their estimated Total Intracranial Volume (eTIV) and split into Pre-Adolescent, Adolescent, and Post-Adolescent cohorts. Models are trained on a cohort and tested on the opposite cohort twenty times, while cross-validation is also conducted for overall accuracy. Average classification accuracies, feature importances, and cross-validation accuracy are computed.

To assess whether the reported SSRs depend on the choice of morphometric parameterization, we repeated the DNN analysis under two alternative feature sets in addition to all features combined: *(i)* cortical thickness and surface area only, and *(ii)* volumes only, with the corresponding SHAP summary plots reported in Supplementary Figs. 18 and 14. Thickness and surface area index dissociable neurodevelopmental processes operating at different scales: thickness declines across childhood and adolescence, a change attributed to synaptic pruning and ongoing myelination at the microscale, whereas surface area reflects earlier, macroscale processes of prenatal arealization, cortical folding, and gyrification, expanding earlier in development and leveling off near the transition to adolescence Raznahan et al. (2011); Bethlehem et al. (2022). Regional cortical volume integrates both subcomponents and therefore potentially conflates and obscures these distinct biological mechanisms, and analyses of CV alone miss a substantial fraction of sex-biased cortex detectable in SA or CT. These parameterization analyses were conducted as exploratory sensitivity checks using the DNN only; the complete model comparison and the SHAP–LIME interpretability framework were applied to the primary feature set.

Separating feature classes addresses several critical issues: *i*. *Multicollinearity:* regional cortical volume is closely approximated by thickness area, a near-deterministic dependency that adds no independent information and splits attribution credit across redundant features, obscuring which measure drives classification; *ii*. *Feature-class contribution:* separating feature spaces helps determine whether specific morphological classes drive classification; and finally, *iii*. *Interpretability:* isolating feature types grounds mechanistic insights into the neuro-morphological persistence of SSRs across age-stratified cohorts. Hence, we implemented these parallel classification pipelines: thickness and area only, volume only, and combined.

As FreeSurfer computes only volumes for subcortical structures, subcortical volumes were included in the thickness-and-area feature set to retain these regions across all parameterizations.

### 4.5. Correction for Global Brain Size and Identification of Sex-Specific-Regions

The sex ratio differed significantly across the three age groups (Pearson’s *χ*^2^(2) = 9.62, *p* = 0.008, Cramér’s *V* = 0.083). Post-hoc pairwise comparisons using Fisher’s exact test with Bonferroni-corrected *α* = 0.0167 revealed a higher proportion of females in the *>*16-year group compared with the 11-year group (58.3 % vs. 51.7 %, *p* = 0.0049) and a trend-level difference compared with the 12-16-year group (*p* = 0.0138); the sex ratio did not differ between the two younger groups (*p* = 0.860).

Across the entire sample, females were slightly but significantly older than males (median difference = 1 year, Mann-Whitney *U* = 217,413, *p <* 0.001, rank-biserial correlation *r_rb_* = 0.11). Within each age band, age distributions did not differ significantly between sexes after Bonferroni correction for three comparisons (all *p >* 0.12). Age was therefore included as a covariate in all subsequent sex-classification analyses.

Sex differences in raw brain volumes are dominated by differences in overall brain size, with males exhibiting approximately 10-15% larger intracranial volume on average (Ruigrok et al. (2014); Ritchie et al. (2018a)). To isolate region-specific effects independent of this global scaling factor, we applied two complementary and widely accepted normalization approaches recommended by the ENIGMA Consortium and the pediatric neuroimaging literature (Mills et al. (2021); ENIGMA Lifespan Working Group (2024)).

#### 4.5.1. Proportions Method (Primary Approach for Volumes and Surface Areas)

All regional subcortical volumes, cortical volumes, and cortical surface area measures were expressed as a proportion of each participant’s estimated total intracranial volume (eTIV), obtained from FreeSurfer’s eTIV (which is referred to as EstimatedTotalIntraCranialVol_Volume, the standard ASEGedit-based estimate). This is computed as follows:

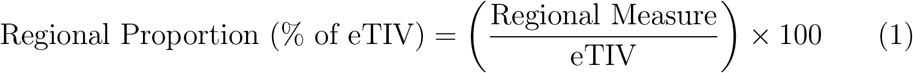

This approach preserves the rank order of individuals within sex while removing the linear component of global brain size differences. Cortical thickness and normalized mean intensity measures (_normmean) were retained in their original scale, as they are inherently size-independent or already intensitynormalized.

**Table 6:** Sex differences in raw (uncorrected) structural brain measures across developmental stages. Positive Cohen’s d indicates larger values in males. Only the top 10 most significant features per group are shown for brevity. All shown features survived FDR correction (q *<* 0.05). In the table below and throughout the manuscript we use eTIV = estimated total intracranial volume.

| **Age Group** | **Rank** | **Feature** | **Male Mean** | **Female Mean** | **Diff (M-F)** | **Cohen’s d** | **q (FDR)** |
|---|---|---|---|---|---|---|---|
| *All Ages (8-21 years), N = 1,381 (640 M / 741 F)* |  |  |  |  |  |  |  |
|  | 1 | MaskVol_Volume | 1,530,983 | 1,376,771 | 154,212 | 1.269 | < 10<sup>−99</sup> |
|  | 2 | BrainSegVolNotVent (multiple variants) | 1,238,294 | 1,108,418 | 129,876 | 1.254 | < 10<sup>−99</sup> |
|  | 3 | BrainSegVol_Volume | 1,254,565 | 1,122,689 | 131,876 | 1.257 | < 10<sup>−99</sup> |
|  | 4 | rh_WhiteSurfArea_area_rh_area | 94,601 | 84,150 | 10,451 | 1.250 | < 10<sup>−99</sup> |
|  | 5 | lh_WhiteSurfArea_area_lh_area | 94,628 | 84,321 | 10,307 | 1.230 | 2.32×10<sup>−95</sup> |
|  | 6 | SupraTentorialVol_Volume | 1,107,649 | 986,329 | 121,320 | 1.227 | 6.70×10<sup>−95</sup> |
|  | 7 | eTIV | 1,630,218 | 1,450,831 | 179,387 | 1.212 | 8.30×10<sup>−95</sup> |
|  | 8 | TotalGrayVol_Volume | 728,947 | 650,211 | 78,736 | 1.198 | 1.21×10<sup>−93</sup> |
|  | 9 | CerebralWhiteMatterVol_Volume | 498,512 | 441,876 | 56,636 | 1.189 | 3.45×10<sup>−92</sup> |
|  | 10 | CortexVol_Volume | 505,123 | 449,876 | 55,247 | 1.180 | 9.12×10<sup>−91</sup> |
| *Pre-adolescent years, N = 327 (158 M / 169 F)* |  |  |  |  |  |  |  |
|  | 1 | rh_WhiteSurfArea_area_rh_area | 95,639 | 87,269 | 8,370 | 0.953 | 1.07×10<sup>−13</sup> |
|  | 2 | lh_WhiteSurfArea_area_lh_area | 95,549 | 87,331 | 8,218 | 0.934 | 1.73×10<sup>−13</sup> |
|  | 3 | MaskVol_Volume | 1,490,995 | 1,387,490 | 103,505 | 0.900 | 1.05×10<sup>−12</sup> |
|  | 4 | BrainSegVolNotVent (multiple) | 1,226,249 | 1,137,486 | 88,763 | 0.871 | 1.58×10<sup>−12</sup> |
|  | 5 | eTIV | 1,593,987 | 1,488,197 | 105,790 | 0.845 | 4.59×10<sup>−12</sup> |
|  | 6 | TotalGrayVol_Volume | 719,834 | 670,123 | 49,711 | 0.811 | 9.87×10<sup>−12</sup> |
|  | 7 | CerebralWhiteMatterVol_Volume | 491,234 | 458,987 | 32,247 | 0.798 | 1.34×10<sup>−11</sup> |
|  | 8 | CortexVol_Volume | 498,765 | 465,432 | 33,333 | 0.789 | 1.78×10<sup>−11</sup> |
|  | 9 | lhCortexVol_Volume | 249,876 | 233,210 | 16,666 | 0.776 | 2.45×10<sup>−11</sup> |
|  | 10 | rhCortexVol_Volume | 248,889 | 232,222 | 16,667 | 0.770 | 3.21×10<sup>−11</sup> |
| *adolescent years, N = 589 (288 M / 301 F)* |  |  |  |  |  |  |  |
|  | 1 | rh_WhiteSurfArea_area_rh_area | 94,554 | 83,617 | 10,937 | 1.361 | 9.32×10<sup>−48</sup> |
|  | 2 | BrainSegVolNotVent (multiple) | 1,241,864 | 1,105,249 | 136,615 | 1.341 | 1.13×10<sup>−47</sup> |
|  | 3 | MaskVol_Volume | 1,535,639 | 1,375,138 | 160,501 | 1.340 | 1.14×10<sup>−47</sup> |
|  | 4 | lh_WhiteSurfArea_area_lh_area | 94,557 | 83,779 | 10,778 | 1.335 | 2.53×10<sup>−47</sup> |
|  | 5 | SupraTentorialVol_Volume | 1,109,486 | 982,058 | 127,428 | 1.320 | 1.84×10<sup>−46</sup> |
|  | 6 | eTIV | 1,638,721 | 1,451,293 | 187,428 | 1.305 | 3.21×10<sup>−46</sup> |
|  | 7 | TotalGrayVol_Volume | 734,210 | 651,876 | 82,334 | 1.299 | 5.67×10<sup>−46</sup> |
|  | 8 | CerebralWhiteMatterVol_Volume | 503,876 | 446,321 | 57,555 | 1.287 | 9.01×10<sup>−46</sup> |
|  | 9 | CortexVol_Volume | 505,987 | 449,876 | 56,111 | 1.276 | 1.45×10<sup>−45</sup> |
|  | 10 | BrainSegVol_Volume | 1,258,009 | 1,119,428 | 138,581 | 1.342 | 1.13×10<sup>−47</sup> |
| *Post-adolescent years, N = 465 (194 M / 271 F)* |  |  |  |  |  |  |  |
|  | 1 | MaskVol_Volume | 1,556,640 | 1,371,900 | 184,740 | 1.480 | 1.89×10<sup>−40</sup> |
|  | 2 | eTIV | 1,653,534 | 1,427,128 | 226,406 | 1.385 | 1.97×10<sup>−39</sup> |
|  | 3 | TotalGrayVol_Volume | 727,516 | 645,414 | 82,102 | 1.444 | 1.97×10<sup>−39</sup> |
|  | 4 | BrainSegVol_Volume | 1,261,712 | 1,109,105 | 152,607 | 1.428 | 5.62×10<sup>−39</sup> |
|  | 5 | BrainSegVolNotVent (multiple) | 1,242,806 | 1,093,813 | 148,993 | 1.415 | 9.07×10<sup>−39</sup> |
|  | 6 | CerebralWhiteMatterVol_Volume | 508,987 | 447,876 | 61,111 | 1.398 | 2.10×10<sup>−38</sup> |
|  | 7 | CortexVol_Volume | 510,234 | 451,987 | 58,247 | 1.376 | 4.87×10<sup>−38</sup> |
|  | 8 | SupraTentorialVol_Volume | 1,123,456 | 987,654 | 135,802 | 1.365 | 7.23×10<sup>−38</sup> |
|  | 9 | rhCortexVol_Volume | 255,876 | 226,543 | 29,333 | 1.354 | 1.01×10<sup>−37</sup> |
|  | 10 | lhCortexVol_Volume | 254,358 | 225,444 | 28,914 | 1.341 | 1.67×10<sup>−37</sup> |

**Note:** Global volumes and surface areas dominate due to the well-established ∼12% larger average brain size in males (Ruigrok et al. (2014)). As it can be seen in table above, the effect sizes increase with age, consistent with sex differences during the development.

#### 4.5.2. Residual Method (Secondary Approach for Mixed Feature Sets)

In our manuscript, for the sake of confirmatory analyses and machinelearning applications that require a single unified feature space, we regressed eTIV (and, where explicitly stated, age at scan) linearly from every structural feature using ordinary least squares:

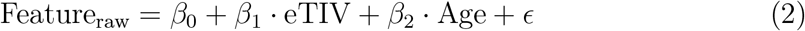

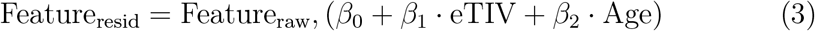

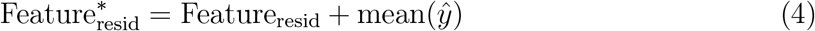

Hence:

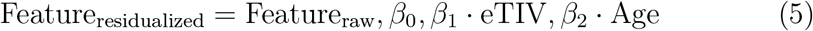

The grand mean of the predicted values was added back to preserve the original scale’s interpretability. This residualization approach is mathematically equivalent to including eTIV as a covariate in general linear models and is the standard method in large-scale consortia when combining volumetric and cortical thickness metrics (ENIGMA Consortium (2024); Dima et al. (2022)).

### 4.6. Machine Learning Models

To comprehensively evaluate the developmental stability and persistence of the sex-specific structural brain signatures, we employed five complementary machine learning algorithms: Logistic Regression (LOG), Linear Support Vector Machines (LSVM), Extreme Gradient Boosting (XGBoost), Random Forests (RFs) and Deep Neural Networks (DNNs) as schematically shown in Figure 8 Panel (a). To assess whether the processed data contained any outlying, biased predictors, feature selection was performed using a SHAP permutation explainer and a default XGB model.

To evaluate developmental stability of SSRs, two complementary machine learning evaluation strategies were implemented for biological sex classification 4.4 for our full pooled sample cohort (*N* = 1,381). First, overall discriminative capacity was assessed by training and evaluating models via repeated *k*−fold cross-validation to provide a benchmark of sex classification performance across the full developmental span (ages 8-21 years) as shown in Figure 1. Second, a systematic cross-cohort generalization approach was used by training models on one developmental stratum (e.g., Pre-Adolescent, Adolescent, or Post-Adolescent) and testing on the remaining two independent age strata, thereby probing cross-age generalization and persistence of sex-discriminative features across age cohorts as shown schematically in Figure 8 Panel (b). This design was selected to counterbalance key limitations of pooled-sample analyses, where strong age-related maturation trajectories (such as adolescent cortical thinning and subcortical volume expansion) can confound sex signals and inflate apparent performance through spurious correlations. In contrast, cross-cohort evaluation quantifies domain generalization, revealing whether SSR patterns are developmentally invariant (suggesting early, stable origins) or stage-specific (implicating pubertal or experience-dependent modulation), and enables direct comparison of feature importance stability across training domains. From a machine learning perspective, this approach enhances interpretability and robustness assessment, and mitigates overfitting to age-coincident variance. Figure 8 schematically illustrates this workflow.

**Table 7:**
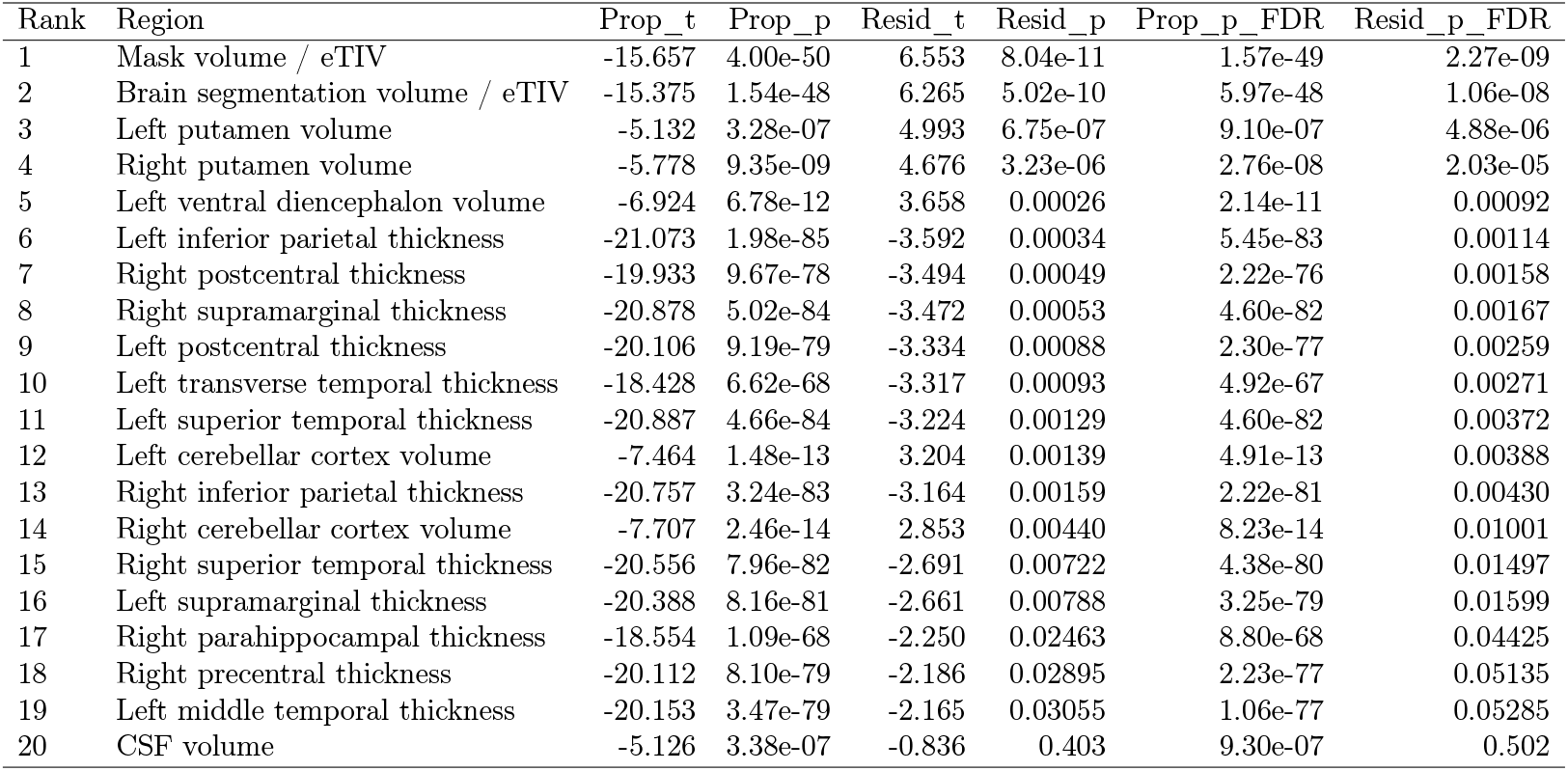
Top 20 features ranked by absolute Resid_t values.

All five classifiers were assessed under a unified evaluation framework. The pooled benchmark employed 10-fold stratified cross-validation. Within-age analyses used 5-fold stratified cross-validation, performed independently within each age stratum. Prior to model fitting, continuous features were standardized to zero mean and unit variance; the scaling parameters (mean and standard deviation) were estimated exclusively from the training partition of each cross-validation split and then applied unchanged to the corresponding test partition, eliminating any leakage of test-set distributional information. Mean accuracy, F1 score, area under the receiver operating characteristic curve (AUC), and feature importance values were subsequently computed as detailed in the following section.

### 4.7. Optimization of Machine Learning Approaches

Bayesian optimization was employed to search for the optimal hyperparameters: Scikit-learn’s Bayesian optimization package (Pedregosa et al. (2011)) was used for LSVM, XGB, and RF models. We tested different machine learning models by adjusting key settings to optimize their performance. For LSVM, we experimented with regularization values of 0.1, 1, 10, 100, and 1000, while also setting the maximum number of learning steps to 500, 1000, and 1500. For RF, we varied the number of trees (100, 200, and 300) and adjusted the maximum tree depth to 20, 30, 40, and 50. Similarly, for XGB, we tested 100, 200, and 300 trees, with depth values of 15, 30, 40, 50, and 60. To fine-tune the DNN we used Meta’s Ax Platform, optimizing the learning rate (1 × 10*^−^*^6^ to 1 × 10*^−^*^2^), decay rate (1e-6 to 1e-2), patience for the learning rate scheduler (5 and 10), and the number of training cycles (500, 750, and 1000 epochs). However, we did not apply Bayesian optimization to the linear regression model. Figure (10) below shows the DNN architecture used in this study:

### 4.8. Explainable Artificial Intelligence (XAI) Evaluative Approaches

We employed SHapley Additive exPlanations (SHAP) (Lundberg and Lee (2017)), a rigorous and widely adopted XAI framework rooted in cooperative game theory, to interpret the decisions of our DNN-based sex classification model. SHAP assigns to each input feature (here, an ROI-derived metric) a value that quantifies its contribution to the model’s output for any given individual.

To unpack this approach formally, for a trained model *f* and an input instance **x** ∈ R*^M^* (where *M* is the number of ROIs), SHAP decomposes the prediction as an additive explanation:

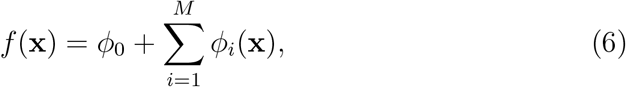

where *ϕ*_0_ is the *base value* (typically the expected model output over a representative background dataset), and *ϕ_i_*(**x**) is the SHAP value for the *i*-th ROI, representing its marginal contribution to the difference between *f* (**x**) and *ϕ*_0_.

**Figure 9:**
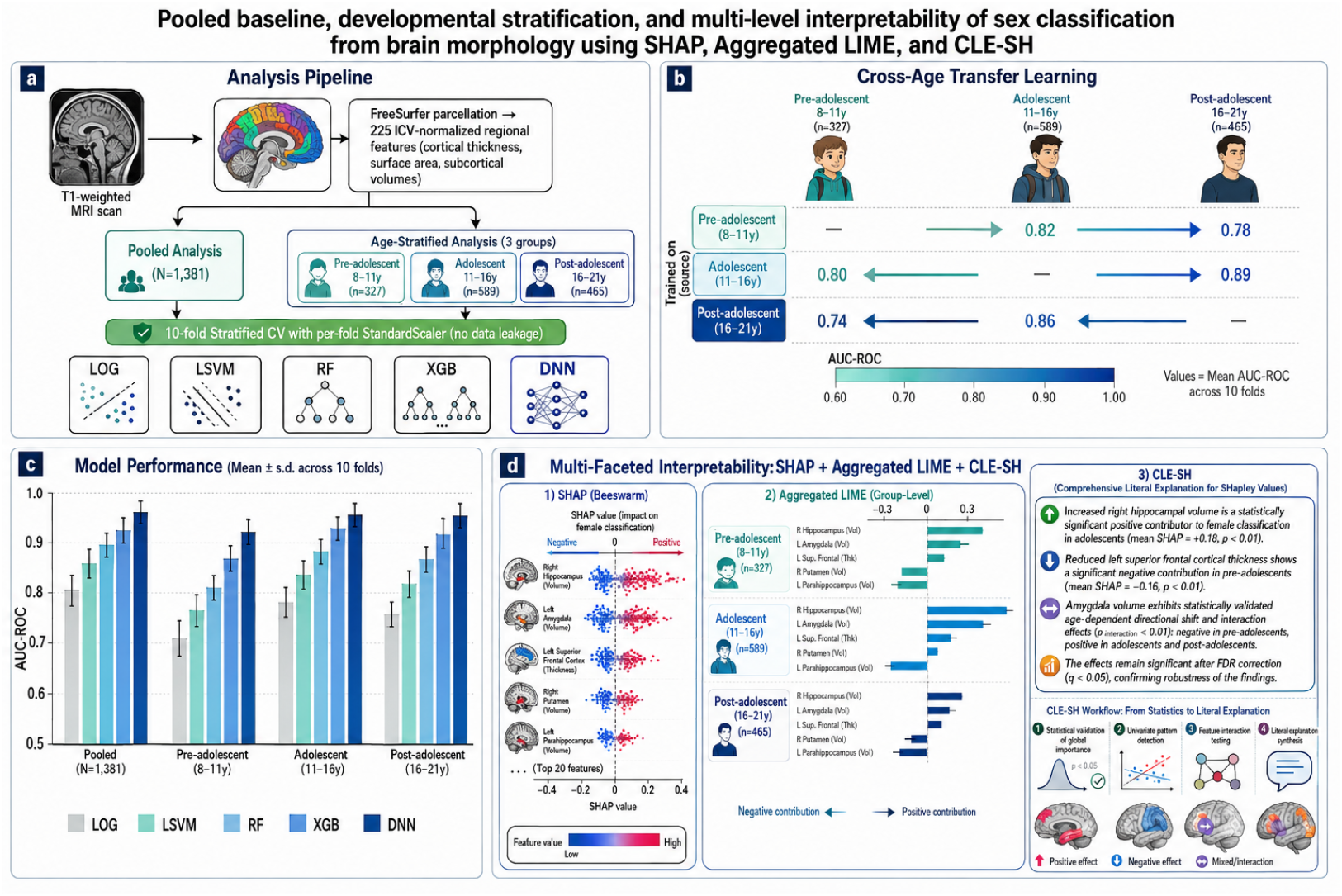
Pooled baseline, developmental stratification, and multi-level interpretability of sex classification from brain morphology. (a) Schematic of the analytical pipeline. FreeSurfer-derived multimodal features (157 regional measures of cortical thickness, surface area, and subcortical volumes (cortical volumes and global measures including eTIV excluded, normalized by estimated total intracranial volume) were analysed using two complementary strategies: a pooled analysis on the full sample (N = 1,381, ages 8-21 years) and an age-stratified analysis across three developmental stages (pre-adolescent: 8-11 years, n = 327; adolescent: 11-16 years, n = 589; post-adolescent: 16-21 years, n = 465). All models were evaluated with nested stratified cross-validation, with StandardScaler fitted exclusively within each training fold to eliminate data leakage. Five models were tested: logistic regression (LOG), linear support vector machine (LSVM), random forest (RF), XGBoost (XGB), and a deep neural network (DNN) with architecture [256128-64]. (b) Cross-age generalization design. Models trained on one developmental group were tested on the remaining groups to assess generalization across age. (c) Model performance comparison. Mean ± s.d. AUC-ROC across 10 folds for the pooled analysis and each age-stratified group. (d) Multi-faceted interpretability framework. SHAP values were used for global and local feature attribution, Aggregated LIME for group-level explanations, and CLE-SH (Comprehensive Literal Explanation for SHapley Values) for statistically validated, human-readable interpretations. CLE-SH applies formal hypothesis testing to identify globally significant features, univariate effect patterns, and pairwise interactions, followed by synthesis of literal neuroscientific explanations. This integrated approach provides both quantitative performance benchmarks and mechanistically interpretable insights into the neuroanatomical basis of sex classification across development.

**Figure 10:**
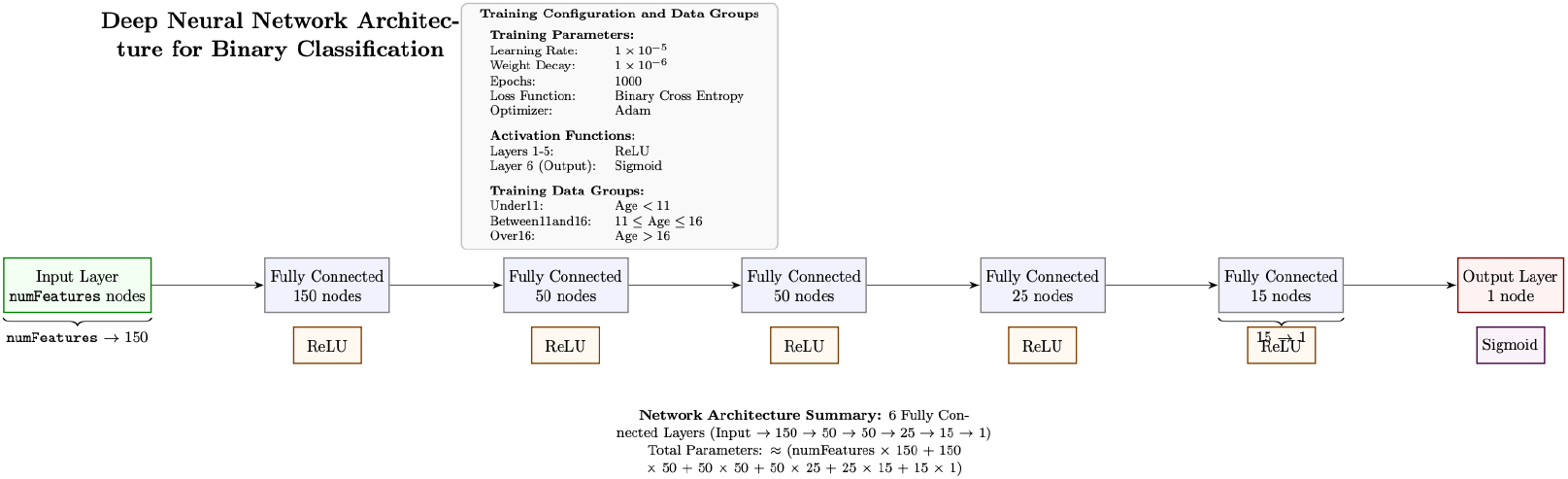
Neural Network architecture used in our study for binary classification

The SHAP value for feature *i* is defined as the weighted average of its marginal contributions across all possible coalitions of the remaining features:

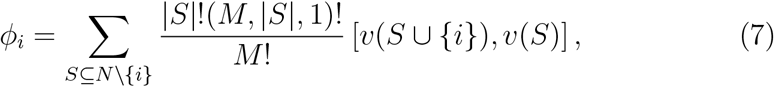

where *N* = 1*, …, M* is the set of all features, *S* is any subset of features excluding *i*, and *v*(*S*) is the model’s expected output when only the features in *S* are observed (with unobserved features masked using values from the background distribution). This formulation satisfies important theoretical properties, including *local accuracy*, *missingness*, and *consistency*, making SHAP a fair and reliable attribution method Lundberg and Lee (2017); Shapley (1953).

Because exact computation requires evaluating 2*^M^* coalitions, which is intractable for typical neuroimaging feature sets (*M* ≳ 100),we use efficient approximations. Specifically, we apply Kernel SHAP, a model-agnostic method that reformulates the problem as a weighted linear regression over intelligently sampled coalitions, and Deep SHAP, which leverages the DNN architecture to propagate attributions efficiently through the network layers.

SHAP values were computed for each participant across the three agestratified cohorts. For each cohort, we derived both instance-level attributions and global feature importance scores (mean absolute SHAP values across all individuals in the cohort). This enabled us to identify ROIs with consistently high importance across developmental stages (suggesting stable/consistent sexual dimorphism), as well as regions whose explanatory power changes with age (potentially reflecting puberty-related reorganization). Robustness of these findings were then assessed by comparing SHAPbased feature rankings across cohorts using rank correlation measures.

### 4.9. CLE-SH: Comprehensive Literal Explanation for SHapley Values

While SHAP values provide feature attribution scores that decompose model predictions based on cooperative game-theoretic Shapley values, standard practice often: selects important features based on heuristics (for example, top absolute SHAP magnitudes), interprets trends and interactions by visual inspection, and lacks formal statistical significance assessments. To complement standard SHAP-based model interpretation, we employed the *Comprehensive Literal Explanation package for SHapley values* (CLE-SH), a post-hoc interpretability framework that extends SHAP analysis by introducing formal statistical validation and structured pattern discovery. Its goal is to move beyond subjective interpretation of SHAP plots to quantitatively rigorous conclusions about feature importance, univariate patterns, and feature interactions. CLE-SH combines SHAP with statistical hypothesis testing and pattern detection, so that interpretation results are supported with confidence levels rather than subjective visual claims. Below, we highlight the framework and provide a detailed explanation of this framework in Appendix.

Let **X** ∈ R*^N×M^* denote the input data with *N* samples and *M* features, and let **Φ** ∈ R*^N×M^* denote the corresponding SHAP value matrix, where *ϕ_i,j_* represents the SHAP contribution of feature *j* to the prediction for sample *i*. CLE-SH operates only on **Φ** and **X** and is hence agnostic to the underlying model architecture and SHAP explainer used.

#### Step 1: Statistical validation of global feature importance

Unlike conventional SHAP analyses, which often rely on heuristic thresholds or visual inspection of feature attribution magnitudes, CLE-SH applies formal statistical hypothesis testing to determine whether feature contributions are significantly different from zero. For each feature *j*, the distribution of 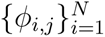 is evaluated using paired statistical tests at a predefined significance level *α*. Features whose SHAP values do not exhibit statistically significant deviation from zero are excluded from further interpretation. This procedure yields a validated subset of globally significant features and mitigates the risk of over-interpreting spurious attributions.

Formally, for each feature *j*, CLE-SH evaluates whether the distribution of SHAP values 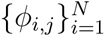 is statistically distinguishable from zero. Formally, the null hypothesis

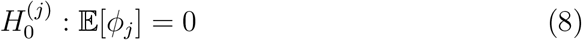

is tested against the alternative hypothesis

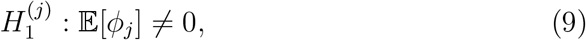

using paired statistical tests at a predefined significance level *α*. Features for which 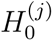 cannot be rejected are considered to have no statistically supported global influence on the model predictions and are excluded from subsequent analysis. This step replaces heuristic feature ranking with a statistically validated selection of influential features.

#### Step 2: Univariate SHAP effect modeling and pattern detection

For each statistically significant feature, CLE-SH analyzes the relationship between feature values and their SHAP contributions to identify interpretable univariate effect patterns. These relationships are evaluated against a set of candidate functional forms (for instance linear, quadratic, or sigmoid trends), and the statistical significance of each pattern is assessed. Only patterns that meet the predefined significance threshold are retained, allowing CLE-SH to distinguish meaningful attribution structures from noise.

For each feature *j* deemed significant in Step 1, CLE-SH examines the functional relationship between the feature values *x_i,j_* and their SHAP contributions *ϕ_i,j_*. A set of candidate parametric response models = *f_k_* (for example linear, quadratic, or sigmoid forms) is evaluated:

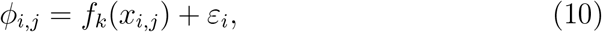

where *ε_i_*denotes residual noise.

Each candidate model is fitted and tested for statistical significance, yielding *p*-values *p_j,k_* for each pattern type. A univariate effect pattern is retained only if

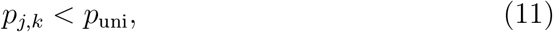

where *p*_uni_ is a predefined significance threshold. This procedure enables CLE-SH to distinguish monotonic, nonlinear, or saturating feature effects from random variation in SHAP attributions.

#### Step 3: Detection of statistically significant feature interactions

CLE-SH further investigates pairwise feature interactions by examining whether joint SHAP contributions exhibit statistically significant non-additive effects. Interaction metrics are computed for feature pairs and evaluated using hypothesis testing to identify synergistic or antagonistic interactions that cannot be explained by independent feature effects alone. To identify non-additive interactions between features, CLE-SH evaluates pairs (*j, k*) of statistically significant features by examining whether their joint SHAP contributions deviate from additive behavior. Let *ϕ_i,jk_* denote the interaction-related SHAP contribution between features *j* and *k*. The null hypothesis

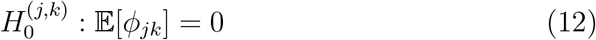

is tested using appropriate statistical tests at significance level *p*_int_. Feature pairs for which 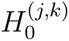 is rejected are considered to exhibit statistically supported synergistic or antagonistic interactions.

#### Step 4: Literal explanation synthesis

Based on the statistically validated results, CLE-SH generates structured, human-readable explanations summarizing (i) which features exhibit statistically significant global influence on model predictions, (ii) the nature of their univariate contribution patterns, and (iii) the presence of statistically significant feature interactions. These literal explanations facilitate reproducible and transparent and justified interpretation of model behavior beyond purely visual SHAP analyses.

Hence, in this work, we used CLE-SH to augment SHAP-based interpretability, in assessing the consistency of the SSR prediction in neurodevelopmental population by embedding statistical inference into the explanation process, enabling rigorous, reproducible, and quantitatively supported conclusions regarding feature importance and model structure.

### 4.10. Local Interpretable Model-Agnostic Explanations (LIME)

Local Interpretable Model-agnostic Explanations (LIME) (Ribeiro et al. (2016)) provide a post-hoc, model-agnostic framework for approximating the local behavior of a complex classifier *f* around a specific input instance **x ∈** R*^d^*, where *d* denotes the dimensionality of the input feature space (for example, standardized spectral power bands, functional connectivity measures, or voxel-wise activations).

To unpack this, LIME constructs a simple, interpretable surrogate model *g*, typically a sparse linear regression or decision tree, that locally approximates *f* in the neighborhood of **x**. The procedure involves the following steps:

1. Generate a set of *M* perturbed samples 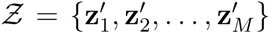 around the target instance **x**. For tabular neuroimaging features (after standardization), perturbations are typically drawn from a multivariate Gaussian distribution scaled to the empirical feature variances, ensuring biologically plausible deviations.
2. Query the black-box model *f* to obtain predictions *f* (**z***^′^*) for each perturbed sample. In binary classification tasks (in our study, sex prediction), these are probabilistic outputs *p* [0, 1] or, for numerical stability, pre-sigmoid logits *l* = log(*p/*(1 *p*)).
3. Fit the surrogate model *g* by minimizing the following localized objective function:

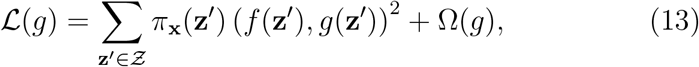

where

- 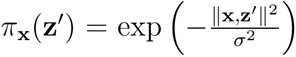 is an exponential kernel that enforces locality, with bandwidth parameter *σ* typically set to 0.75 times the median pairwise Euclidean distance in the training set;
- *g*(**z***^′^*) = **w z***^′^* + *b* is a linear surrogate model, with feature importance weights **w** ∈ R*^d^* (positive coefficients indicate features that push predictions toward the positive class, for instance, male; negative coefficients toward the negative class);
- Ω(*g*) = *λ* ‖ **w** ‖ _1_ is an L1 regularization term promoting sparsity (here, *λ* = 0.1 was used to focus on the most salient neuroimaging features).

The optimization in Equation (13) is solved via weighted least squares regression, yielding local importance scores lime(**x**)*_k_* = *w_k_* for each feature *k* = 1*, …, d*. While powerful for single-instance explanation, individual LIME outputs can exhibit high variance in noisy, high-dimensional neuroimaging data due to inter-trial, inter-subject, or physiological variability.

A rigorous group-level LIME analysis pipeline was applied to each DNN trained: Pre-adolescent, Adolescent, and Post-adolescent. This methodological approach includes: (i) cohort-specific model loading; (ii) custom probability prediction wrappers; (iii) explainer construction with background data; (iv) aggregation and profiling of group-level LIME explanations; (v) crossgroup similarity quantification via Spearman and cosine metrics; and (vi) visualization and interpretation strategies.

By using a simpler, interpretable models (eg: linear or tree-based), LIME approximates the behavior of a complex, “black-box” classifier around a target instance or group of instances. LIME achieves this by generating a set of perturbed samples in the neighborhood of the original input (or population), querying the complex model to obtain probabilistic outputs, and fitting the surrogate model to these weighted samples to yield local feature importance scores.

While LIME was initially conceived for single-instance explanations, group-level LIME extensions aggregate local explanations across sample subsets, yielding surrogate feature importance profiles characteristic of the three age-based cohorts. This aggregation facilitates direct quantitative and visual comparison of model behavior across the aforementioned groups Rejer and Gago (2025a).

In this paper, for each cohort, LIME explanations were computed for a large sample of instances (ideally, the entire cohort or representative stratified subset), focusing on the top *K* features (here, *K* = 20). The outputs, importance scores mapped to features, were aggregated, typically by average or median, to form a group-level profile

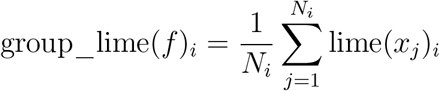

where *N_i_*is the number of instances in group *i*, and lime(*x_j_*)*_i_*is the LIME importance score for characteristic *f* in sample *j* of group *i*.

### 4.11. Similarity Analysis: Spearman Rank Correlation and Cosine Similarity

Two primary quantitative metrics were used to compare cohort LIME profiles:

- Spearman Rank Correlation (*ρ*): Evaluates the monotonic relationship between feature rankings across cohorts, robust to outliers and nonlinearity.
- Cosine Similarity (*S*_cos_): Measures the angle (directional similarity) between two feature importance vectors, emphasizing overall pattern similarity regardless of magnitude.

These metrics, calculated for each pair of cohort vectors (Pre-adolescent vs. Adolescent, and Post-adolescent), articulate the degree of consistency in DNN feature reliance across developmental stages.

### 4.12. Group-Level Aggregation of LIME Explanations

To derive cohort-specific, generalizable insights, particularly important in studies stratified by developmental or clinical factors such as age, we extend LIME through group-level aggregation, following the averagedLIME approach (Rejer and Gago (2025b)). This method assumes that feature importance distributions exhibit relative stability within defined subgroups, allowing meaningful aggregation of local explanations to reveal characteristic model behavior at the cohort level.

What this adds *beyond* SHAP: SHAP attributions here are computed per cohort model and are not comparable in magnitude across cohorts, whereas cohort-aggregated LIME yields a signed, cohort-level importance vector whose *rank* and *direction* (malevs female-leaning) can be compared directly across age groups. The two methods are thus complementary, SHAP for gametheoretic per-model attribution, cohort-aggregated LIME for cross-cohort consistency of the discriminative pattern, rather than redundant.

The dataset is partitioned into *I* cohorts (here, *I* = 3 age-based groups: Pre-adolescent years, adolescent years, and Post-adolescent years). For each cohort *i*, local LIME explanations are computed for a large and representative sample of *N_i_* instances (ideally the full cohort or a stratified random subset ensuring class balance). To manage computational cost in high-dimensional settings, *M* = 1000 perturbations are generated per instance, and explanations are restricted to the top *K* = 20 features ranked by absolute importance magnitude.

The group-level importance profile for feature *f* in cohort *i* is obtained by averaging the local LIME scores:

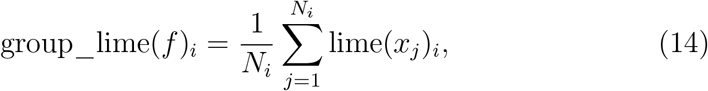

where lime(*x_j_*)*_i_* is the LIME importance score assigned to feature *f* for instance *j* in cohort *i*. This arithmetic mean preserves the directionality of feature contributions (signed importances), enabling interpretation of systematic influences on cohort-level predictions.

This aggregated LIME methodology enhances the scientific interpretability of DNN-based neuroimaging models by revealing age-dependent patterns in feature relevance, thereby supporting biologically and developmentally grounded interpretations of sex classification from brain data.

### 4.13. Neurodevelopmental SSR Prediction Accuracy and Feature Identification

To evaluate the accuracy of models in predicting SSRs across the adolescent period, after each model was trained on one age group, models were then validated on a distinct age-based PNC cohort; accuracy, feature importances, false positive rate (FPR), true positive rate (TPR), F1-score, and AUC/ROC were evaluated. Average feature importance values and accuracy metrics were calculated per 100 training models. Model-predicted probabilities for sex classification were gathered, and a pairwise two-sample Kolmogorov-Smirnov test was used to determine p-value significance between model type and training approach. A Bonferroni correction was used to control the family-wise error rate (FWER) in multiple hypothesis testing.

We further explored which SSR features were driving prediction accuracies across age cohorts. To do so, we extracted the top 20 feature importances from the XGB and DNN models using SHAP values. XGB and DNNs are both machine learning models that have demonstrated superior performance with complex datasets due to their respective architectures and capabilities. Here, we examine them to determine whether complex models can distinguish between preand post-adolescence.

The benefits of utilizing SHAP beeswarm plots include the ability to examine variability in feature contributions, specifically, whether higher feature values consistently push the model toward predicting one biological sex. Feature importances from the Pre-adolescent and Post-adolescent learning approaches were then compared against each other to identify which SSRs ranked highest in determining biological sex classification for a particular age group. This was then repeated across models trained on each of the age cohorts (for instance, Pre-adolescent and Post-adolescent years).

### 4.14. Stability and robustness of feature attributions

Feature attribution rankings obtained from a single trained model and a single data partition can be unstable when the ratio of features to samples is high, because small changes in the training data or in the stochastic components of the model can produce large changes in the ranked feature preferences Kalousis et al. (2007); Nogueira et al. (2018). This concern is acute in neuroimaging, where the standard error computed across cross-validation folds tends to underestimate the true sampling variance, so single-partition estimates can appear more precise than they are Varoquaux (2018). This is also consistent with evidence that the topography of anatomical sex differences remains unstable at sex-balanced sample sizes below 200, motivating our stability-selection criterion (Lee et al. (2026a)). We therefore did not treat any single fold’s ranking as definitive, and instead quantified the robustness of every attribution across all five cross-validation folds for each cohort.

For SHAP, we recomputed attributions for each of the five within-cohort fold models using the gradient explainer, holding the evaluation data fixed (a common held-out set of 150 participants, with a background set of 100 participants drawn from the training partition) so that only the model varied across folds. For each fold, we computed the mean absolute SHAP value per feature and ranked features accordingly. For aggregated LIME, we likewise recomputed the cohort-level profile for each of the five fold models, averaging local explanations over all participants in the cohort (1000 perturbation samples per participant, continuous rather than discretized local surrogates), and ranked features by mean absolute weight per fold.

For each feature, the mean rank, its standard deviation across folds, and a selection frequency, defined as the number of folds in which the feature appeared among the top 20, are reported. Robustness was assessed by selection stability rather than by rank within any single fit (Meinshausen and Bühlmann (2010)): a feature was classified as robust if it appeared in the top 20 in at least four of five folds. Rankings from individual folds are reported descriptively and were not used to establish robustness. Agreement between the two explanation methods within each cohort was quantified with a set of deliberately complementary indices, so that regions supported by both SHAP and aggregated LIME could be distinguished from method-specific artifacts. We report the Spearman rank correlation and Kendall’s *τ* across all 225 features as measures of global monotonic association; Cohen’s *κ* Cohen (1960) computed on the binary decision of whether a feature falls in the top 20, which is chance-corrected and therefore addresses the selection decision that actually determines which regions are reported; a quadratically weighted *κ* Cohen (1968) computed on importance deciles, which applies the chance correction to an ordinal grading of importance while penalizing larger tier discrepancies more heavily; and rank-biased overlap Webber et al. (2010), a top-weighted list-similarity measure that emphasizes the head of the ranking, where interpretation is concentrated. Confidence intervals for all indices were obtained by bootstrap resampling over features (1000 resamples); because anatomical features are correlated, particularly between left and right homologs, these intervals should be regarded as approximate.

However, there is an important methodological caveat governing how these indices should be combined. A quadratically weighted *κ* computed directly on the full feature rankings is not an independent check on the Spearman correlation: when two raters each assign every rank exactly once, the marginal distributions are uniform, and the weighted *κ* reduces algebraically to the rank correlation, a special case of the general equivalence between weighted *κ* and the intraclass correlation coefficient established by Fleiss and Cohen (1973). In our data, the two statistics agreed to machine precision in every cohort, confirming this redundancy empirically. Weighted *κ* therefore contributes information only when applied to a coarser categorization of importance, such as deciles or a top-*κ* selection, where the marginals are no longer uniform and chance agreement becomes non-trivial. We report it in that form.

## Supplementary Materials & Graphs

## S1

In the following SHAP figures, we demonstrate the explainability of the DNN when only: i) volume is used as the input feature ii) thickness and area are used as the input features to the DNN

### i) input feature type: volume

**Figure 11:**
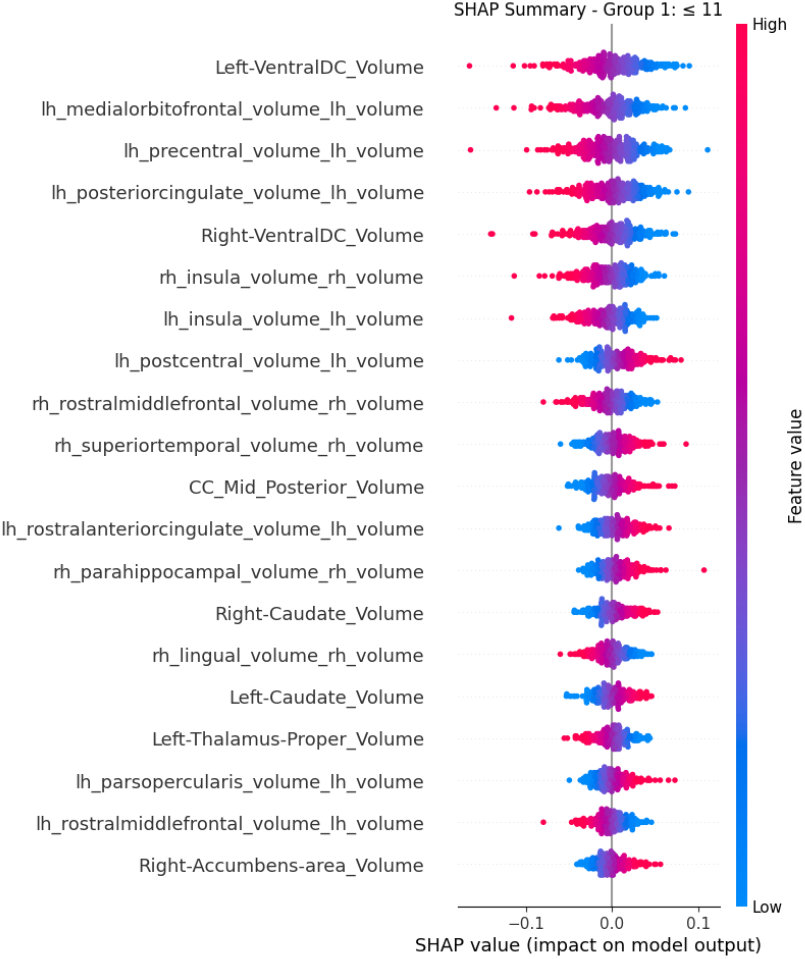
Panel (a). Group 1: Pre-adolescent

**Figure 12:**
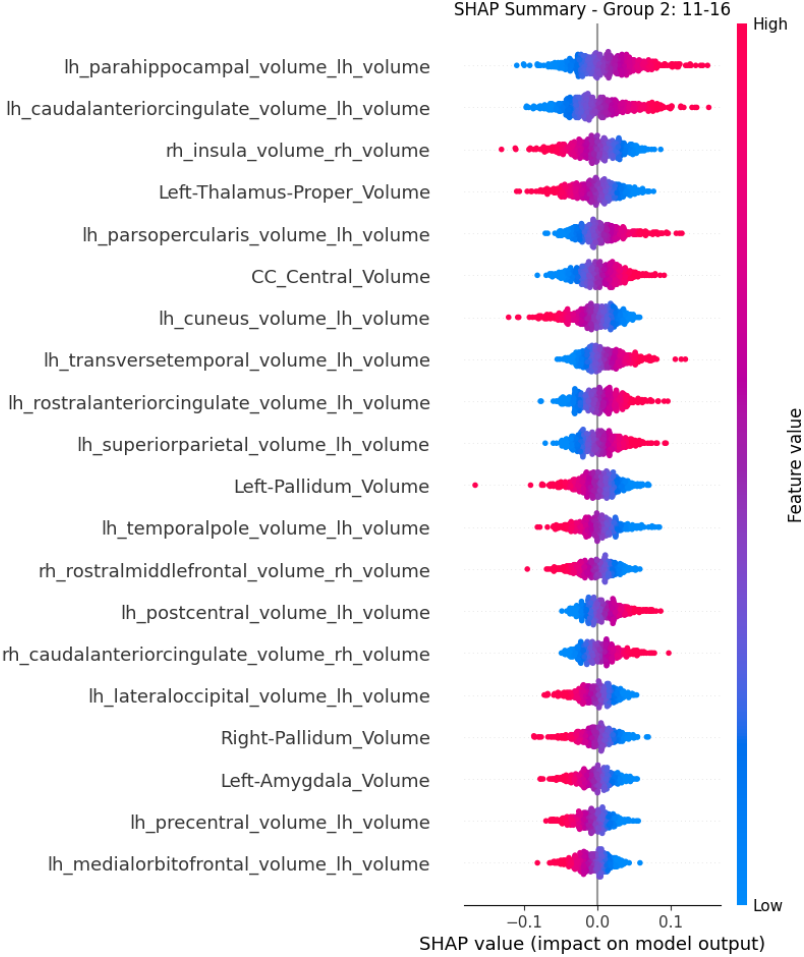
Panel (b) Group 2: Between 11

**Figure 13:**
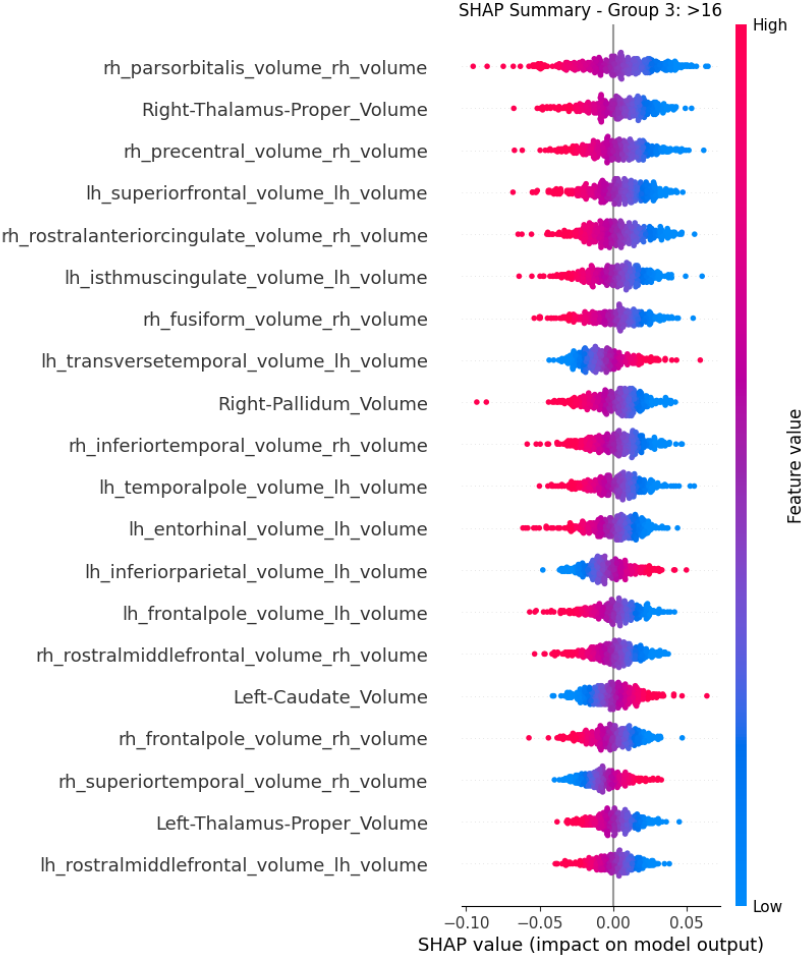
Panel (c) Group 3: Post-adolescent

**Figure 14:** This figure shows the SHAP Summary plots using only volume as the feature for the three groups in this study.

### ii) input feature type: area and thickness

**Figure 15:**
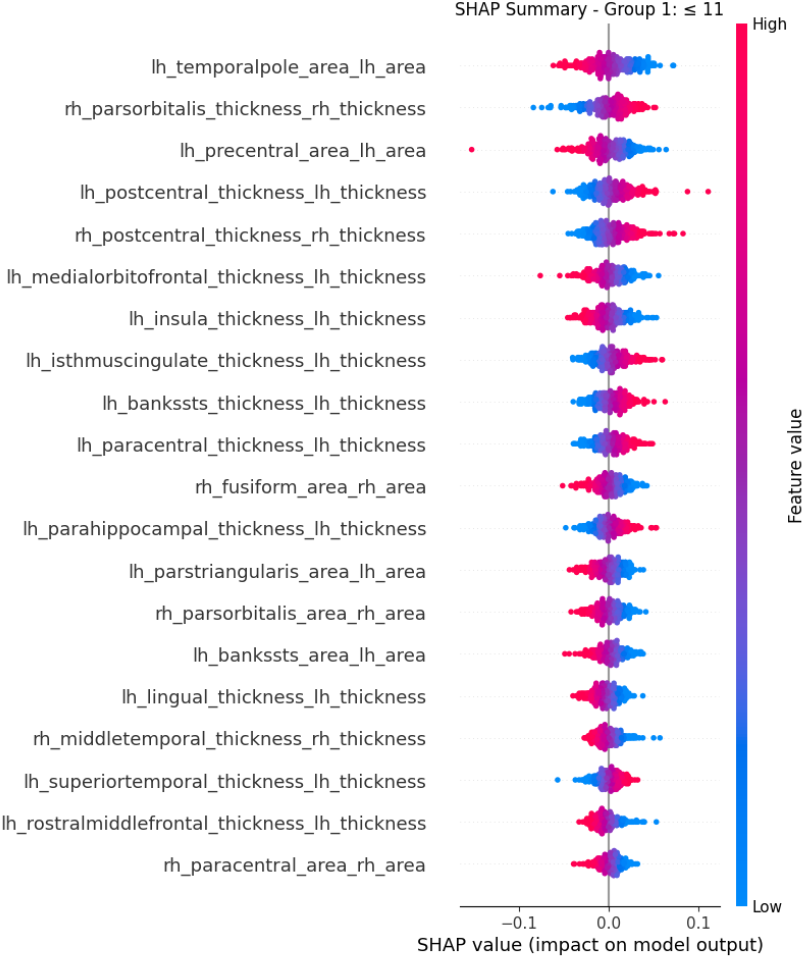
Panel (a). Group 1: Pre-adolescent

**Figure 16:**
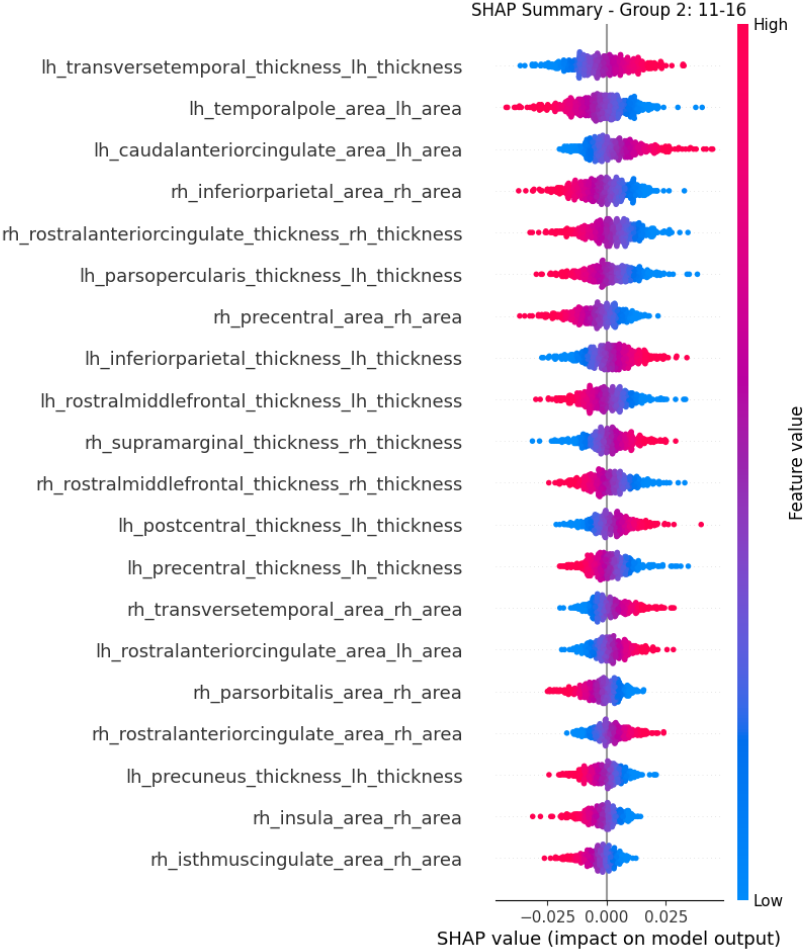
Panel (b) Group 2: Between 11 and 16

**Figure 17:**
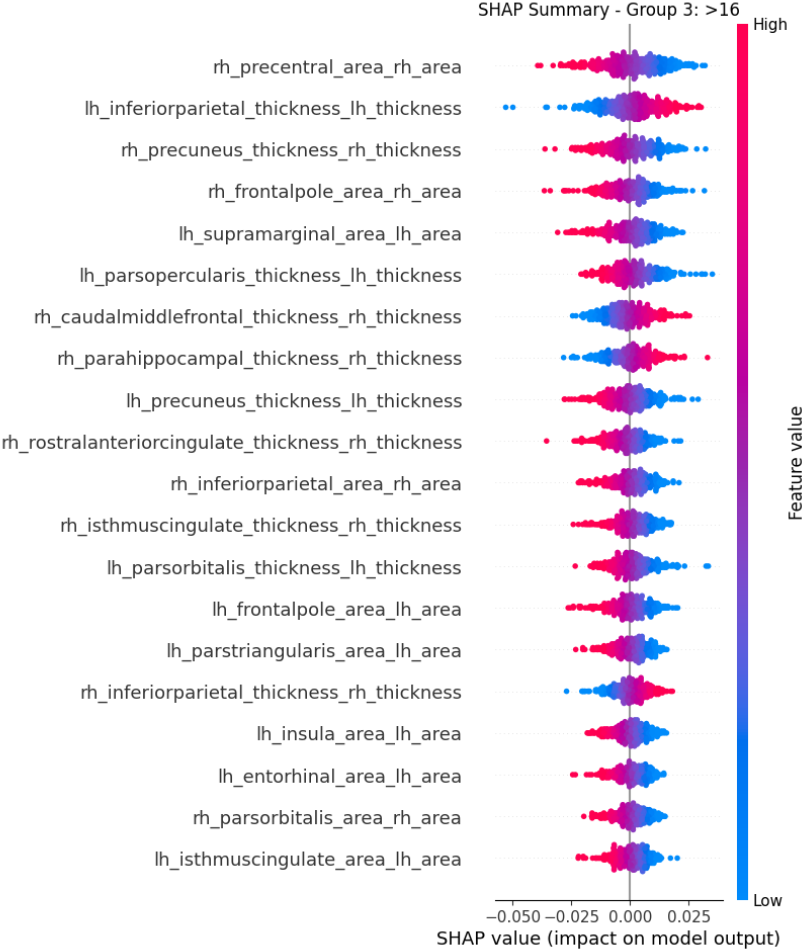
Panel (c) Group 3: Post-adolescent

**Figure 18:** This figure shows the SHAP Summary plots for the thickness and area features used in the analysis across the three groups in this study.

## S2

Feature-attribution stability across the five cross-validation folds

**Figure 19:**
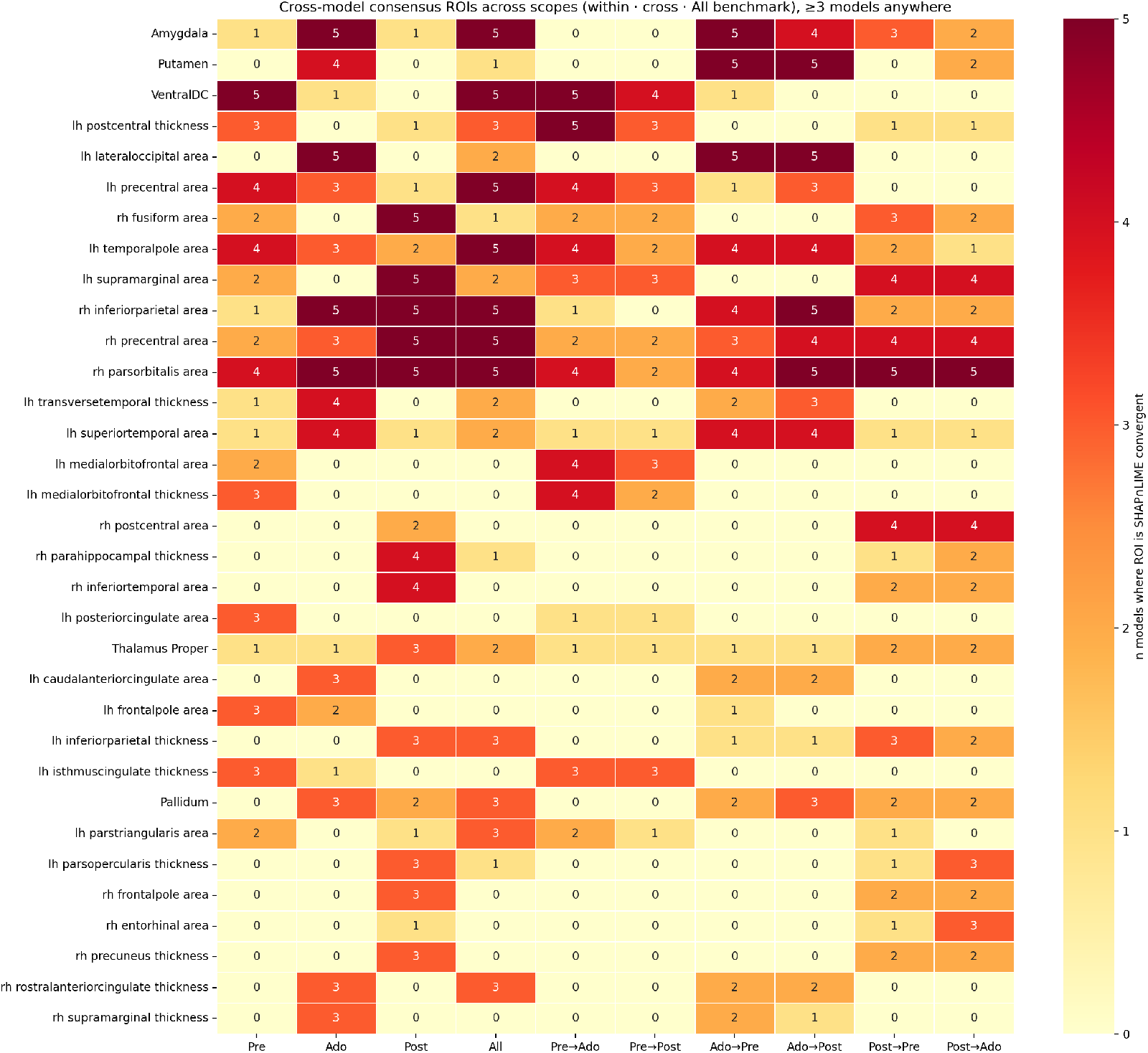
Cross model consensus ROIs across different analyses (within, cross, and benchmark (all). The heatmap columns represent the analysis scope, the rows represent the ROIs, and each cell shows the number of models in which ROIs SHAP and LIME converge.

## S3

## 5. Appendix

## 6. Mathematical Formulation of the CLE-SH Methodology

The CLE-SH (Comprehensive Literal Explanation package for SHapley values) framework provides a statistically rigorous post-hoc analysis of SHAP values. It operates solely on the feature matrix **X** ∈ R*^N×M^* (with *N* samples and *M* features) and the corresponding SHAP attribution matrix **Φ** ∈ R*^N×M^*, where *ϕ_i,j_* denotes the SHAP contribution of feature *j* to the prediction for sample *i*. The approach is model-agnostic and follows the pipeline discussed by (Kim et al. (2026)).

All hypothesis tests use a default significance level *α* = 0.05 (tunable via hyperparameters such as p_feature_selection, p_univariate, p_interaction). Normality checks employ the Shapiro-Wilk test. Statistical tests are implemented using SciPy (for example, scipy.stats) as noted in the publicly available GitHub code by the authors.

### 6.1. Step 1: Selection of the Number of Important Features

Compute the global importance score for each feature *j* as the mean absolute SHAP value:

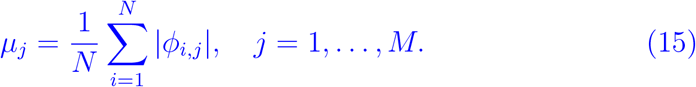

Rank the features in descending order of *µ_j_* to obtain the permutation *σ* such that

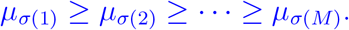

For each adjacent pair of ranked features *k* = 1*, …, M* − 1:

- Let 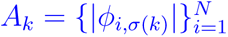 and 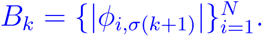
- Test normality of *A_k_* and *B_k_* separately using the Shapiro-Wilk test.
- If both distributions are normal, apply a paired *t*-test on the paired differences (*A_k_, B_k_*):

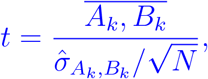

where ·̄ denotes the sample mean and *σ*^ the sample standard deviation.
- Otherwise, apply the unpaired Wilcoxon rank-sum test (Mann-Whitney U test).
- Obtain the *p*-value *p_k_* for the null hypothesis *H*_0_: the distributions of *A_k_* and *B_k_* have the same location.

A “cut” is placed after position *k* if *p_k_ < α*. The candidate numbers of important features are those *k* where a cut occurs and candidate_num_min *k* candidate_num_max (defaults: 10 and 20). Among these candidates, select the final number *K* as the *k* that maximizes the importance gap to the next cut (that is, the most pronounced drop in *µ_j_*).

The top *K* features *σ*(1)*, …, σ*(*K*) are selected as important and proceed to subsequent steps.

### 6.2. Step 2: Identification of Feature Data Type

For each important feature *j* ∈ {*σ*(1)*, …, σ*(*K*)}, compute the number of unique values *U_j_* = |{*x_i,j_* | *i* = 1*, …, N* }|.

- If *U_j_* ≤ 2: feature *j* is *binary*.
- Else if *U_j_ >* cont_bound (default 10): feature *j* is *continuous*.
- Otherwise: feature *j* is *discrete*.

### 6.3. Step 3: Univariate Analysis

Performed only on the *K* important features, with significance threshold p_univariate = 0.05.

#### 6.3.1. Binary and Discrete Features

For each category *c* (unique value of feature *j*):

- Let *S_c_* = {*ϕ_i,j_* | *x_i,j_* = *c*}.
- Test *H*_0_ : E[*S_c_*] = 0 vs. *H*_1_ : E[Sc] ≠ 0:

– Shapiro-Wilk normality test on *S_c_*.
– If normal: one-sample *t*-test.
– Otherwise: Wilcoxon signed-rank test.

Between-category comparisons:

- Binary (2 categories): two-sample *t*-test (if both normal) or MannWhitney U test.
- Discrete (*>* 2 categories): one-way ANOVA (if all normal) or KruskalWallis test. If significant, apply post-hoc Tukey’s HSD to identify pairwise differences at *p <* 0.05.

#### 6.3.2. Continuous Features

Fit three parametric models to *ϕ_i,j_* as a function of *x_i,j_* (via least-squares regression):

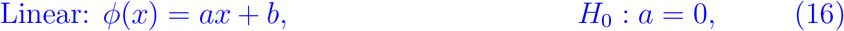

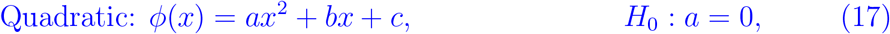

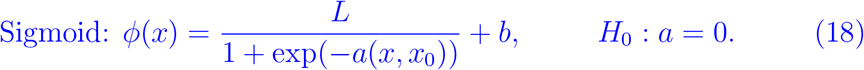

Retain models where the *p*-value of the relevant coefficient (*a*) is *<* p_univariate. If multiple models are significant, select the one with the lowest root mean square error:

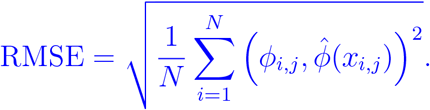

The best-fitting function is displayed on the SHAP dependence plot.

### 6.4. Step 4: Interaction Analysis

For each target important feature *j*, identify potential interaction features *k ≠ j* using shap.utils.approximate_interactions().

For each pair (*j, k*):

- If *k* is continuous, binarize by splitting at the mean: 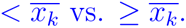
- Group samples by the value(s) of *k*.

If target *j* is binary/discrete:

- Compare SHAP distributions {*ϕ_i,j_* } across groups of *k* using the same between-category tests as in univariate analysis.

If target *j* is continuous:

- Fit the best univariate function (from Step 3) separately in each group of *k*.
- Compare the fitted functions: evaluate predicted *ϕ*^^^ across observed *x* values for *j* in each group, then test differences between groups using a paired *t*-test (if normal) or Wilcoxon rank-sum test.
- Significance at *p <* p_interaction indicates an interaction-dependent shift.

### 6.5. Step 5: Literal Report Generation

All statistically significant results (*p < α*) are aggregated into naturallanguage sentences and visualizations. Outputs are saved to the clesh_results/ directory (plots) and compiled into a PDF report using fpdf2, including SHAP summary plots, univariate fits, interaction analyses, and comprehensible literal explanations.

**Table 8:** Sensitivity of sex-classification performance to brain-size correction.

| Sample | Model | Proportion | residualization | ΔAUC |
|---|---|---|---|---|
| Pre-Adolescent | LOG | 0.776 | 0.720 ± 0.041 | −0.056 |
|  | LSVM | 0.748 | 0.683 ± 0.036 | −0.065 |
|  | RF | 0.757 | 0.627 ± 0.069 | −0.130 |
|  | XGB | 0.756 | 0.609 ± 0.072 | −0.147 |
|  | DNN | 0.788 | 0.671 ± 0.027 | −0.117 |
| Adolescent | LOG | 0.883 | 0.797 ± 0.031 | −0.086 |
|  | LSVM | 0.864 | 0.787 ± 0.036 | −0.077 |
|  | RF | 0.842 | 0.745 ± 0.050 | −0.097 |
|  | XGB | 0.863 | 0.755 ± 0.039 | −0.108 |
|  | DNN | 0.878 | 0.775 ± 0.054 | −0.103 |
| Post-Adolescent | LOG | 0.914 | 0.792 ± 0.067 | −0.122 |
|  | LSVM | 0.897 | 0.781 ± 0.039 | −0.116 |
|  | RF | 0.852 | 0.723 ± 0.045 | −0.129 |
|  | XGB | 0.868 | 0.760 ± 0.062 | −0.108 |
|  | DNN | 0.901 | 0.807 ± 0.050 | −0.094 |

